# Monocytes shape the neuroprotective and immunomodulatory effects of mesenchymal stromal cell-derived extracellular vesicles

**DOI:** 10.64898/2026.06.01.727369

**Authors:** Chen Wang, Yiqiao Zhang, Tobias Tertel, Yanis Mouloud, Xiaolong Liu, Nina Hagemann, Ayan Mohamud Yusuf, Janine Gronewold, Jan-Kolja Strecker, Aurel Popa-Wagner, Matthias Mack, Jens Minnerup, Matthias Gunzer, Bernd Giebel, Dirk M. Hermann

**Affiliations:** Department of Neurology, University Hospital Essen, University of Duisburg-Essen, Essen, Germany; Center for Translational and Behavioral Neurosciences, University Hospital Essen, University of Duisburg-Essen, Essen, Germany; Institute of Transfusion Medicine, University Hospital Essen, University of Duisburg-Essen, Essen, Germany; Department of Neurology, University of Lübeck and University Hospital Schleswig-Holstein, Campus Lübeck, Lübeck Germany; Department of Neurology, University of Münster, Münster, Germany; Center of Normal and Pathological Ageing, University of Medicine and Pharmacy Craiova, Craiova, Romania; Department of Nephrology, University Hospital Regensburg, Regensburg, Germany; Institute of Experimental Immunology and Imaging, University Hospital Essen, University of Duisburg-Essen, Essen, Germany

**Keywords:** Ischemic stroke, exosome, antiinflammation, M2-like monocyte/macrophage, non-classical monocyte, neutrophil, T cell

## Abstract

**BACKGROUND:** Mesenchymal stromal cell–derived extracellular vesicles (MSC-EVs) exert neuroprotective effects in ischemic stroke largely through immunomodulatory mechanisms. Monocytes are first-line responders to MSC-EVs. Their contribution to MSC-EV-induced neuroprotection remains poorly understood. This study investigated the role of monocytes in shaping neuroprotective responses to MSC-EVs after ischemic stroke.

**METHODS:** Male C57BL/6J mice were exposed to transient middle cerebral artery occlusion (MCAO). Monocytes were depleted using pharmacological (clodronate liposomes), immunological (anti-CCR2), or genetic (*Mrp8-Cre*^+/–^ *Nr4a1*^fl/fl^) approaches removing total, CCR2^+^, or Ly6C^low^ monocytes, respectively. In additional cohorts, neutrophils and T cells were simultaneously depleted by anti-Ly6G or anti-CD4/CD8 antibodies. Small EVs from clonally expanded immortalized MSCs were administered intravenously. Neurological deficits, ischemic injury, and immune responses were analyzed up to 72 hours post-MCAO. Complementary *ex vivo* studies were performed, in which MSC-EVs were administered to monocyte-depleted or non-depleted peripheral blood mononuclear cells (PBMCs) obtained from acute ischemic stroke patients.

**RESULTS:** In ischemic mice with intact monocyte compartment, MSC-EVs reduced neurological deficits, infarct volume, neuronal injury, and brain leukocyte infiltrates. These protective effects were abolished in monocyte-depleted mice, particularly following CCR2^+^ monocyte depletion. Under these conditions, MSC-EV treatment exacerbated neurological deficits, ischemic injury, and leukocyte infiltration, accompanied by neutrophil and T cell expansion and overactivation. Depletion of neutrophils or T cells prevented the EV-induced worsening of stroke outcome in monocyte-deficient mice. Ly6C^low^ monocytes played a crucial role in orchestrating immune responses to MSC-EVs. Their depletion abolished EV-induced neuroprotection. In stroke patient PBMCs, MSC-EVs induced phenotypic reprogramming of monocytes, whereas they promoted CD4^+^ and CD8^+^ T cell activation in the absence of monocytes.

**CONCLUSIONS:** Monocytes shape the immunomodulatory actions of MSC-EVs. In their absence, MSC-EVs trigger neutrophil and T cell overactivation that worsens stroke outcome. These findings highlight the importance of monocyte- and T cell-related potency assays for the clinical translation of MSC-EV therapies.

## Introduction

Reperfusion therapies (i.e., intravenous thrombolysis and endovascular thrombectomy) considerably enhanced clinical outcome of ischemic stroke patients in recent years.^1^ Despite these benefits, the majority of patients still exhibit long-term disability, and secondary brain injury driven by post-ischemic inflammation plays a significant role in chronic stroke impairments.^2^ There is a lack of treatments in the clinical setting able to reverse this inflammatory brain injury. Mesenchymal stromal cell (MSC)-derived small extracellular vesicles (EVs) have emerged as promising candidates for stroke therapy, which bring together the advantages of the pleiotropic mode-of-action of cell therapies with simple handling of EV samples and the lack of malignant transformation risks.^3,4^ In mouse and rat transient middle cerebral artery occlusion (MCAO) models, we and others have shown that intravenously administered MSC-EVs potently promote neurological recovery, reduce infarct volume, and enhance neuronal survival and plasticity.^5-8^ Considering their potent actions, MSC-EVs are rapidly approaching clinical proof-of-concept trials.^3,9^

The recovery-promoting effects of MSC-EVs in ischemic stroke models decisively depend on their immunomodulatory properties. In MCAO mice, MSC-EVs with neuroprotective activity effectively reduced the brain infiltration of leukocytes, namely of polymorphonuclear neutrophils, monocytes/ macrophages, and lymphocytes.^5,7,10^ In MCAO mice, in which neutrophils were depleted by a monoclonal anti-Ly6G antibody immediately after reperfusion onset, MSC-EV treatment did not reduce neurological deficits and ischemic brain injury.^7^ Likewise, MSC-EVs failed to induce vascular remodeling and angiogenesis in mice with delayed neutrophil depletion starting one day after MCAO.^10^ These studies underline the importance of immunomodulatory effects for mediating both the neuroprotective and neurorestorative actions of MSC-EVs.

Monocytes are pivotal targets for MSC-EVs in a wide set of inflammatory and degenerative pathologies.^11,12^ When fluorescently labeled MSC-EVs are added to peripheral blood mononuclear cells (PBMCs), they are preferentially internalized by monocytes and macrophages rather than other leukocytes.^13,14^ MSC-EVs have potent effects on monocytes/ macrophages, promoting their polarization from a proinflammatory (M1-like) toward an antiinflammatory (M2-like) phenotype, as shown in mouse myocardial infarction models.^15,16^ While classical Ly6C^high^ monocytes are rapidly recruited to injured tissue closely associated with inflammatory injury where they may convert from M1-like to M2-like cells, non-classical Ly6C^low^ patrolling monocytes contribute to the resolution of inflammation, tissue homeostasis and brain repair.^17,18^ We previously observed that MSC-EVs significantly reduce monocyte accumulation in the ischemic brains of MCAO mice.^7,19^ However, whether monocytes are required for the neuroprotective effects of MSC-EVs, has not been investigated in ischemic stroke studies.

To evaluate the role of monocytes in mediating MSC-EV responses, we herein performed transient MCAO studies, in which we depleted monocytes by pharmacological, immunological or genetic tools, and intravenously administered small EVs obtained from clonally expanded immortalized MSCs (ciMSCs), which we generated by a modified human telomerase reverse transcriptase (hTERT) strategy.^20^ We have previously shown in a mouse model of neonatal hypoxia–ischemia that small EVs from ciMSCs exert neuroprotective effects comparable to primary MSC-EVs.^21,22^ We now explored whether the neuroprotective effects of ciMSC-EVs (hereafter referred to as MSC-EVs) after MCAO depend on monocytes. To this end, we explored the effect of MSC-EVs in MCAO mice, in which monocytes, neutrophils and/or T cells were depleted and used a PBMC assay using blood samples of acute ischemic stroke patients to mimic effects of MSC-EVs under clinically relevant conditions *ex vivo*.

## Materials and Methods

### Legal issues, randomization and statistical planning

A detailed description of all methods is provided in the Supplemental Material. Experiments were performed with local government approval (State Office for Consumer Protection and Food North Rhine-Westphalia, Recklinghausen) in accordance to EU (Directive 2010/63/EU) and local institutional guidelines for the care and use of laboratory animals in accordance with ARRIVE guidelines for reporting animal experiments.^23^ Experiments were strictly randomized and blinded. Statistical planning assumed an alpha error of 5% and a beta error (1–statistical power) of 20%.

### MSC culture, EV preparation, and characterization

The MSC-EV preparation used in the present study was identical to our previous studies.^21,22^ Briefly, ciMSCs were generated from our human MSC stock (source 41.5) derived from bone marrow aspirates of healthy donors as previously described.^21,22^ Small EVs were harvested from ciMSC conditioned media using an optimized polyethylene glycol 6000 precipitation protocol followed by ultracentrifugation, as previously reported.^7,19,21^ MSC-EVs were characterized in accordance with the Minimal Information for Studies of Extracellular Vesicles 2018 (MISEV2018) guidelines,^24^ including nanoparticle tracking analysis (NTA), bicinchoninic acid (BCA) assay, and imaging flow cytometry for EV surface marker expression.

### Depletion of monocytes, neutrophils, and T cells

For depletion of peripheral blood monocytes/ macrophages, control liposomes or clodronate-loaded liposomes (LIPOSOMA, Amsterdam, Netherlands) were injected intravenously 24 hours before (50 mg/kg) and 24 and 48 hours after (30 mg/kg) MCAO.^25,26^ For monocyte depletion, 20 μg of IgG isotype antibody (rat IgG2b; BE0090; BioXCell, West Lebanon, NH; as control) or monoclonal anti-CCR2 antibody MC-21 (kindly provided by Matthias Mack) were injected intravenously 12 hours before and 24 and 48 hours after MCAO.^27^ For abolishing non-classical Ly6C^low^ monocytes, which patrol along vascular endothelium and maintain brain tissue homeostasis,^18^ *Mrp8-Cre*^+/–^ mice on a C57BL/6J background were crossbred with *Nr4a1*^fl/fl^ mice on a C57BL/6J background generating *Mrp8-Cre*^+/–^ *Nr4a1*^fl/fl^ mice (provided by Jens Minnerup). In defined subgroups, neutrophils were depleted by intraperitoneal injection of 200 μg of anti-Ly6G antibody (BE0075-1; BioXCell) 24 hours before and 24 hours after MCAO,^7^ or CD4 and CD8 T cells were depleted by the intraperitoneal injection of anti-CD4 (BE0119; BioXCell) and anti-CD8 (BE0061; BioXCell) antibody 24 hours before (100 µg each) and 24 hours after (50 µg each) MCAO.

### Focal cerebral ischemia and MSC-EV treatment

Focal cerebral ischemia was induced by transient intraluminal MCAO as previously described.^7,19^ Briefly, 10-12-week-old male C57BL/6J mice (Harlan Laboratories, Darmstadt, Germany) or *Mrp8-Cre*^+/–^ *Nr4a1*^fl/fl^ mice on a C57BL/6J background were anesthetized with 1.5% isoflurane (30% O_2_, remainder N_2_O). Rectal temperature was maintained between 36.5 and 37.0 °C using a feedback-controlled heating system (Fluovac, Harvard apparatus, Holliston, MA, U.S.A.). Cerebral blood flow was recorded by laser Doppler flowmetry (Perimed, Stockholm, Sweden) above the core of the middle cerebral artery territory. A silicon-coated 7.0 nylon monofilament (Doccol Corporation, Sharon, MA, U.S.A.) was introduced through a small incision into the common carotid artery and advanced to the carotid bifurcation for MCAO. After 30 min, reperfusion was initiated by monofilament removal. Immediately thereafter, 200 μl normal saline (as vehicle) or 2x10^6^ cell equivalents of MSC-EV preparations dissolved in 200 μl normal saline were administered through the animals’ tail veins.

### Analysis of neurological deficits, infarct volume and brain edema

Neurological deficits were evaluated at 24, 48, and 72 hours after MCAO using the Clark score as previously described.^7,19^ 20-μm-thick coronal brain cryostat sections collected at 1 mm intervals across the forebrain were stained with cresyl violet. In all sections, infarct area was determined using the indirect method that corrects for brain swelling by subtracting area of healthy tissue of the ischemic hemisphere from that of the contralesional hemisphere using Image J software (National Institute of Health, Bethesda, MD, U.S.A.). Infarct volume was determined by integrating infarct areas from all brain levels, and brain edema was measured as ratio of ipsilateral to contralateral hemisphere volume.

### Immunohistochemistry

20-μm-thick coronal brain sections obtained from the rostrocaudal level of the bregma (i.e., the core of the middle cerebral artery territory) were immunolabeled for NeuN (neuronal marker), CD45 (leukocyte marker), Ly6G (neutrophil marker), CD3 (T cell marker), CD31 (vascular endothelium marker), ICAM-1 (intercellular adhesion molecule-1, an inflammation marker on endothelial cells), collagen-IV (a marker of ischemic microvessels), GPIbα (glycoprotein Ibα, a platelet marker), or extravasated serum IgG (a blood-brain barrier permeability marker). Nuclei were counterstained with Hoechst 33342. NeuN stainings were costained with terminal deoxynucleotidyl transferase dUTP nick end labeling (TUNEL) (12156792910; Roche Diagnostics, Mannheim, Germany). Injured NeuN^+^/ TUNEL^+^ neurons, ICAM-1^+^ microvessels, CD45^+^ leukocytes, Ly6G^+^ neutrophils, CD3^+^ T cells, GP-Ibα ^+^ microvascular thrombi, and IgG extravasation were evaluated in the ischemic striatum by counting cell numbers, determining cell densities, or measuring optical densities.

### Flow cytometry of leukocytes

Single cell suspensions for flow cytometry analysis were prepared as previously described.^7,19^ Single cell suspensions were stained with antibody cocktails listed in Supplemental Table S2 and analyzed by a CytoFLEX flow cytometer (Beckman-Coulter) using Kaluza software V2.2 (Beckman-Coulter).

### Peripheral blood mononuclear cell (PBMC) assay

PBMCs were isolated from peripheral blood samples of patients with acute ischemic stroke using a Ficoll gradient. A total of 6□×□10^5^ viable, intact or monocyte-depleted PBMCs were seeded per well in 96-well U-bottom plates (SARSTEDT, Nümbrecht, Germany) in a final volume of 200 µl. Cells were cultured for 3 days in the presence or absence of MSC-EVs (5x10^4^ cell equivalents dissolved in 5 µl). After culture, cells were harvested, stained with fluorochrome-conjugated antibodies (Table S2), and analyzed by flow cytometry with CytoFLEX (Beckman Coulter).

### Statistical analysis

Statistical analysis was performed using GraphPad Prism (version 10.4.1; GraphPad Software, San Diego, California U.S.A.). Normal distribution was assessed in all datasets using the Shapiro-Wilk normality test. Normally distributed data were analyzed by one-way ANOVA followed by Tukey’s post hoc test (comparisons between ≥ 3 groups) or by unpaired or paired two-tailed Student’s t test (comparisons between 2 groups). Non-normally distributed data were evaluated by Kruskal-Wallis test followed by Dunn’s multiple comparisons test (comparisons between ≥ 3 groups) or two-tailed Mann-Whitney U test (comparisons between 2 groups). Data were presented as box plots with median ± interquartile ranges with minimum and maximum values shown as whiskers and individual data points as dots. P<0.05 was considered statistically significant.

## Results

### Monocyte/ macrophage depletion by clodronate liposomes turns neuroprotection by small MSC-EVs into exacerbation of ischemic brain injury

To explore the effect of monocyte/ macrophage depletion on neuroprotective responses to MSC-EVs, we first administered PBS-loaded liposomes (as control) or clodronate liposomes to mice 24 hours before transient MCAO (Figure 1A). Flow cytometric analysis of blood samples confirmed efficient depletion of monocytes 24 hours after intravenous clodronate liposome administration, including all major monocyte subsets— classical Ly6C^high^ inflammatory, Ly6C^int^ intermediate, and non-classical Ly6C^low^ patrolling monocytes (Figure 1B, C). In addition, clodronate liposomes reduced the frequencies of macrophages and NK cells in the blood, while increasing immature and aged neutrophil, CD4^+^ T cell, activated CD8^+^ T cell, and B cell frequencies (Figure S1). Notably, dendritic cell frequencies were not influenced by clodronate liposomes (Figure S1). In MCAO mice treated with PBS-loaded control liposomes, MSC-EV administration immediately after reperfusion significantly reduced neurological deficits over 72 hours post-MCAO compared with vehicle, as assessed by the Clark score, which evaluates both general and focal neurological impairments (Figure 1D). Consistently, infarct volume, brain edema and the density of DNA-fragmented TUNEL^+^/NeuN^+^ injured neurons were significantly reduced by MSC-EVs (Figure 1E, F). These findings confirm that the neuroprotective potency of the ciMSC-EVs we used is equivalent to that of primary MSC-EVs administered earlier by our group.^5,7,10,19^ Notably, monocyte depletion by clodronate liposomes mimicked the neuroprotective effects of MSC-EVs, significantly reducing neurological deficits, infarct volume, brain edema, and neuronal injury (Figure 1D-F). In contrast, in monocyte-depleted mice, MSC-EV treatment not only failed to induce neuroprotection compared with vehicle treatment, but significantly exacerbated neurological deficits, infarct volume, and neuronal injury (Figure 1D-F) and increased the expression of ICAM-1, a leukocyte adhesion molecule, on cerebral microvessels (Figure 1G). MSC-EVs and monocyte depletion did not influence the density of GPIbα ^+^ microvascular thrombi (Figure S2A) or IgG extravasation, a marker of blood-brain barrier (BBB) permeability (Figure S2B), in the previously ischemic striatum, neither when applied alone or as combination treatment. Together, these data indicate that monocytes are essential mediators of MSC-EV-induced neuroprotection, whereas ischemic injury is exacerbated by MSC-EVs in the absence of monocytes.

**Figure 1.**
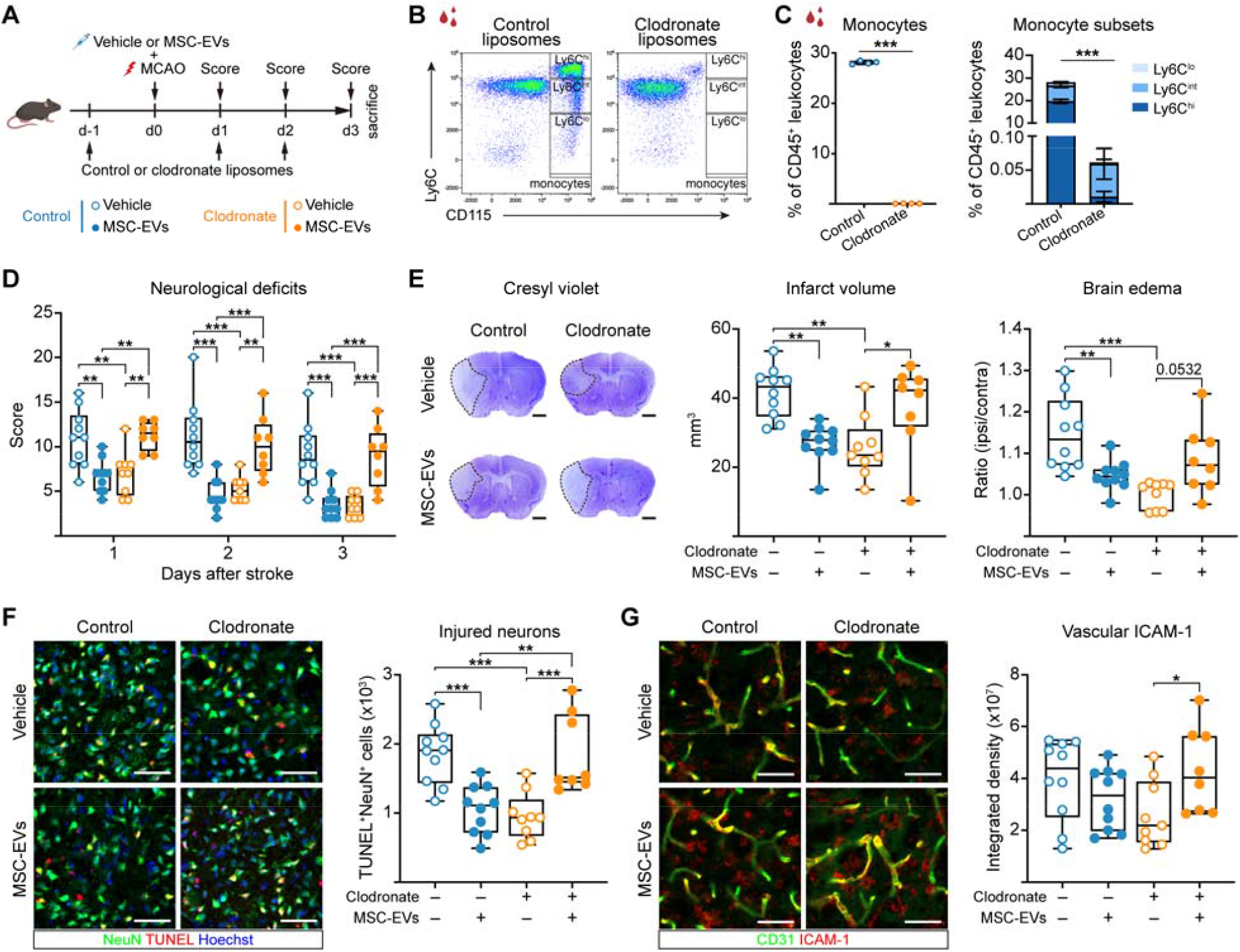
Mesenchymal stromal cell (MSC)-derived extracellular vesicle (EV)-induced neuroprotection turns into exacerbation of ischemic brain injury in monocyte-depleted mice. (**A**) Experimental design: Control or clodronate liposomes were intravenously administered to C57BL/6J wildtype mice 24 hours prior to MCAO (50 mg/kg) and 24 and 48 hours post-MCAO (30 mg/kg each) for monocyte depletion. Vehicle or small EVs obtained from clonally expanded immortalized MSCs (2x10^6^ cell equivalents) were intravenously applied immediately after reperfusion. Neurological deficits were evaluated at 24, 48, and 72 hours after MCAO using the Clark score, followed by animal sacrifice at 72 hours (created with https://BioRender.com). (**B**) Gating strategy for circulating monocytes (CD11b^+^ CD115^+^) in the blood classified as classical Ly6C^high^, intermediate Ly6C^int^, and non-classical Ly6C^low^ monocytes. (**C**) Quantification of monocyte subsets in blood samples of non-ischemic mice 24 hours after intravenous administration of control liposomes or clodronate liposomes. (**D**) Neurological deficits using the Clark score assessed at 24, 48, and 72 hours post-MCAO. (**E**) Infarct volume and hemisphere brain edema evaluated by cresyl violet staining, (**F**) neuronal injury in the ischemic striatum assessed by terminal transferase-mediated dUTP-nick end labeling (TUNEL)/ NeuN labeling, and (**G**) ICAM-1 expression on CD31^+^ microvessels in the ischemic striatum at 72 hours post-MCAO. Representative cresyl violet stainings (in **E**) and immunohistochemistry images (in (**F, G**)) are shown. Data were evaluated by unpaired t test (in (**C**)) or one-way ANOVA with Tukey post hoc test (in (**D-G**)). *p<0.05, **p<0.01, ***p<0.001 (n=4 mice/group (in (**C**)), n=8-10 mice/group (in (**D-G**))). Scale bars: 1 mm (in (**E**)), 50 μm (in (**F, G**)).

### Monocyte depletion converts antiinflammatory activity of MSC-EVs into proinflammatory activity

The upregulation of microvascular ICAM-1 expression by MSC-EVs in monocyte-depleted mice (Figure 1G) suggested facilitated immune cell infiltration into the ischemic brain. We therefore investigated the immunomodulatory effects of MSC-EVs in the presence and absence of monocytes. Ischemic brain, blood, and spleen samples were collected at 3 days after MCAO and analyzed by flow cytometry (Figure 2A). In ischemic brains, monocyte depletion alone by clodronate liposomes significantly reduced total brain-infiltrating CD45^+^ leukocyte numbers, while MSC-EV treatment alone had no overall effect. In contrast, combined monocyte depletion and MSC-EV administration significantly increased brain leukocyte infiltrates (Figure 2B). Analysis of neutrophils within the brain-infiltrating leukocyte population revealed that monocyte depletion and MSC-EV treatment alone significantly reduced brain neutrophil accumulation. Monocyte depletion furthermore decreased the number of activated neutrophils, resting neutrophils, N1-like neutrophils, and N2-like neutrophils in the ischemic brain (Figure 2C, D). Conversely, combined monocyte depletion and MSC-EV treatment tended to increase total neutrophil number (p=0.0733) and significantly increased activated neutrophils in the ischemic brain (Figure 2D). Analysis of lymphocyte subsets showed that monocyte depletion alone significantly reduced post-ischemic brain T cell, activated CD8^+^ T cell, B cell, and NK cell infiltrates (Figure 2E, F, Figure S3A). In contrast, MSC-EVs significantly increased brain T cell, CD4^+^ T cell, CD8^+^ T cell, activated CD8^+^ T cell, and B cell infiltrates in monocyte-depleted mice (Figure 2F). In the blood, neither monocyte depletion nor MSC-EVs alone significantly influenced leukocyte responses compared with vehicle-treated stroke mice (Figure 2G, Figure S3B). However, combined monocyte depletion and MSC-EV treatment significantly increased total leukocyte number, which comprised T cells, CD4^+^ T cells, CD8^+^ T cells, neutrophils, activated neutrophils, aged neutrophils, and immature neutrophils (Figure 2G, Figure S3B). In the spleen, both monocyte depletion alone and combined monocyte depletion and MSC-EV treatment significantly decreased NK cell number (Figure S3C). Flow cytometry of T cell suppressive and exhausted phenotypes revealed that in monocyte-depleted mice treated with MSC-EVs, the number of Tregs, PD-1^+^ CD4^+^ T cells, and PD-1^+^ CD8^+^ T cells was increased in the blood, while PD-1^+^ CD8^+^ T cells were increased in the brain (Figure S4A-C). No significant effects were observed in the spleen (Figure S4D). These findings indicate that monocyte depletion converts the antiinflammatory effects of MSC-EVs into proinflammatory effects, accompanied by the appearance of immunosuppressive and exhausted immune cell phenotypes. Collectively, these data underscore a critical role of monocytes in MSC-EV-induced immunomodulation.

**Figure 2.**
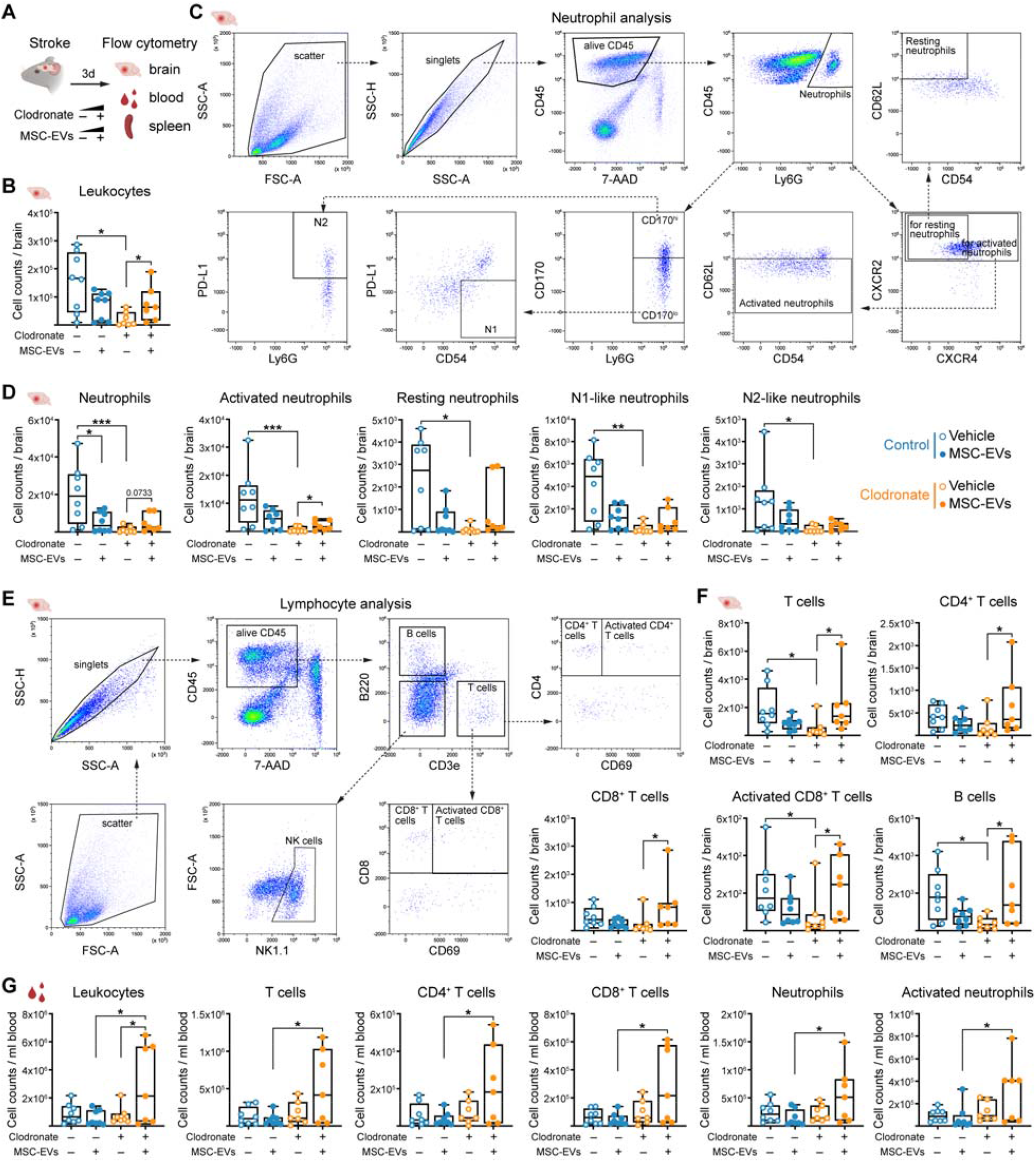
MSC-EVs augment post-ischemic cerebral inflammatory responses and peripheral immune activation in monocyte-depleted mice. (**A**) Experimental design: Mice were intravenously treated with control or clodronate liposomes for monocyte depletion (see Figure 1A), and vehicle or MSC-EVs (2x10^6^ cell equivalents) were intravenously applied immediately after MCAO. Ischemic brain, blood, and spleen samples were collected at 72 hours after MCAO for flow cytometry (created with https://BioRender.com). (**B**) Quantification of total CD45^+^ leukocytes in ischemic brain. (**C**) Gating strategy for analysis of neutrophils within brain-invading leukocytes. (**D**) Quantification of neutrophil subsets in ischemic brain, including neutrophils (Ly6G^+^), activated neutrophils (Ly6G^+^ CXCR2^+^ CD62L^low^), resting neutrophils (Ly6G^+^ CXCR2^+^ CXCR4^-^ CD62L^high^ CD54^low^), N1-like neutrophils (Ly6G^+^ CD170^low^ CD54^+^ PD-L1^−^), and N2-like neutrophils (Ly6G^+^ CD170^high^ PD-L1^+^). (**E**) Gating strategy for lymphocyte subsets. (**F**) Quantification of lymphocyte subsets in ischemic brain, including T cells (CD3e^+^), CD4^+^ T cells (CD3e^+^ CD4^+^), CD8^+^ T cells (CD3e^+^ CD8^+^), activated CD8^+^ T cells (CD3e^+^ CD8^+^ CD69^+^), and B cells (CD3e^−^ B220^+^). (**G**) Quantification of leukocyte subsets in peripheral blood, including total CD45^+^ leukocytes, T cells, CD4^+^ T cells, CD8^+^ T cells, neutrophils, and activated neutrophils. Data were compared by Kruskal-Wallis test with Dunn’s post hoc test or one-way ANOVA with Tukey post hoc test. *p<0.05, **p<0.01, ***p<0.001 (n=7-8 mice/group).

### CCR2^+^ monocytes are indispensable for MSC-EV-induced neuroprotection and antiinflammation after ischemic stroke

Cell depletion by clodronate liposomes is not specific to monocytes/ macrophages. Indeed, other phagocytes, such as dendritic cells, have also been reported to be eliminated by clodronate.^28^ In addition, clodronate liposomes may induce antiinflammatory effects via myeloid cell, specifically neutrophil stunning.^29^ Although we did not observe effects of clodronate liposomes on dendritic cells (Figure S1) and evidence of neutrophil stunning (Figure S1, S3), we next specifically depleted monocytes using the anti-CCR2 antibody MC-21 in MCAO mice to elucidate the role of CCR2^+^ monocyte subsets (Figure 3A). Flow cytometry of blood samples confirmed the depletion of CCR2^+^ classical monocytes after MC-21 delivery (Figure 3B, C), while frequencies of immature neutrophils and activated CD4^+^ and CD8^+^ T cells were increased (Figure S5). In isotype control mice, MSC-EV administration reproducibly reduced neurological deficits, infarct volume, and neuronal injury, when administered immediately after reperfusion (Figure 3D-F). CCR2^+^ monocyte depletion alone reduced neurological deficits, but did not significantly influence infarct volume, brain edema, or neuronal injury (Figure 3D-F). Notably, MSC-EVs again increased neurological deficits, infarct volume, neuronal injury, and ICAM-1 expression in monocyte-depleted mice (Figure 3D-G). Microvascular thrombosis and BBB permeability were unaffected across treatment groups (Figure S6). Flow cytometry revealed that MSC-EVs significantly increased the brain infiltration of leukocytes, particularly neutrophils, activated neutrophils, N1-like neutrophils, and B cells in the ischemic brain of CCR2^+^ monocyte-depleted mice (Figure 3H, I). Treg frequency (percentage of Treg in CD4^+^ T cells) was increased, with a trend toward increased PD-1^+^ Tregs (p=0.0712; Figure S7A). MSC-EVs modestly increased Treg frequency in the blood (Figure S7B) and increased aged neutrophils and Treg frequency in the spleen (Figure S7C). Thus, MSC-EV-induced worsening of stroke outcome in CCR2^+^ monocyte-depleted mice was again associated with exacerbated neutrophil activation, T cell suppression and exhaustion, suggestive of an endogenous protective mechanism activating a compensatory negative feedback loop. These data confirm that CCR2^+^ monocytes are required for the tuning of inflammatory responses following MSC-EV treatment.

**Figure 3.**
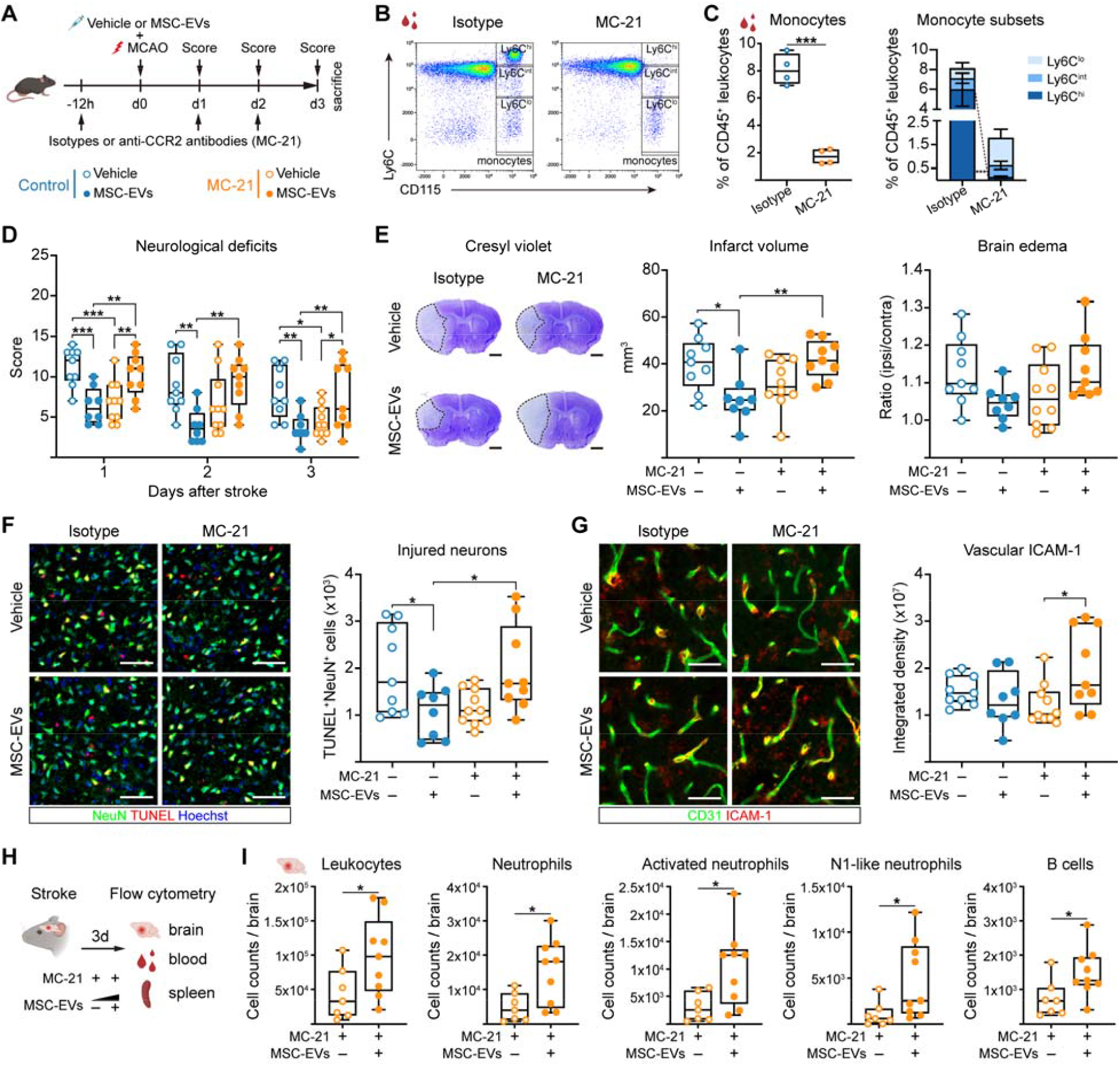
MSC-EVs aggravate ischemic stroke outcome and increase brain inflammatory responses in mice exhibiting CCR2^+^ monocyte depletion. (**A**) Experimental design: Isotype control antibody or monoclonal anti-CCR2 antibody (MC-21) was intravenously injected 12 hours prior to MCAO and 24 and 48 hours after MCAO (20 µg each) for monocyte depletion. Vehicle or MSC-EVs (2x10^6^ cell equivalents) were intravenously administered immediately after reperfusion. Neurological deficits were evaluated at 24, 48 and 72 hours. Animals were sacrificed at 72 hours post-MCAO for histochemical analyses (created with https://BioRender.com). (**B**) Gating strategy for circulating monocytes (CD11b^+^ CD115^+^) in blood classified as classical Ly6C^high^, intermediate Ly6C^int^, and non-classical Ly6C^low^ monocytes. (**C**) Quantification of monocyte subsets in non-ischemic mice 12 hours after the administration of isotype control or MC-21 antibody. (**D**) Neurological deficits assessed at 24, 48, and 72 hours after MCAO using the Clark score. (**E**) Infarct volume and brain edema evaluated by cresyl violet staining, (**F**) neuronal injury in the ischemic striatum assessed by TUNEL/NeuN immunolabeling, and (**G**) ICAM-1 expression on CD31^+^ microvessels in the ischemic striatum at 72 hours post-MCAO. (**H**) Experimental design: Anti-CCR2 antibody (MC-21) were injected for monocyte depletion (as above), and vehicle or MSC-EVs (as above) were intravenously applied immediately after reperfusion. Ischemic brain, blood, and spleen samples were harvested at 72 hours post-MCAO for flow cytometry (created with https://BioRender.com). (**I**) Quantification of leukocyte subsets in ischemic brain, including total leukocytes, neutrophils, activated neutrophils, N1-like neutrophils, and B cells. Representative cresyl violet stainings (in (**E**)) and immunohistochemistry images (in (**F, G**)) are shown. Data were compared by unpaired t test (in (**C, I**)) or one-way ANOVA with Tukey post hoc test (in (**D**-**G**)). *p<0.05, **p<0.01, ***p<0.001 (n=4 mice/group (in (**C**)), n=8-10 mice/group (in (**D**-**G**)), n=7-9 mice/group (in (**I**)). Scale bars: 1 mm (in (**E**)), 50 μm (in (**F, G**)).

### Critical role of non-classical Ly6C^low^ monocytes for post-ischemic neuroprotection and antiinflammation by MSC-EVs

Considering that monocytes were found to regulate immunomodulatory responses, we next investigated the role of non-classical Ly6C^low^ monocyte subsets, which patrol along vascular endothelium and control brain tissue homeostasis^18^ using Ly6C^low^ monocyte-deficient *Mrp8-Cre*^+/–^ *Nr4a1*^fl/fl^ mice (Figure 4A). In a steady state of these mice, circulating Ly6C^low^ monocytes were significantly reduced, whereas Ly6C^high^ and Ly6C^int^ monocyte populations, total macrophage, total neutrophil, and dendritic cell frequencies were unaffected (Figure 4B, C, Figure S8). Moreover, *Mrp8-Cre*^+/–^ *Nr4a1*^fl/fl^ mice exhibited decreased frequencies of M2-like macrophages and resting neutrophils, while immature and aged neutrophil and NK cell frequencies were increased (Figure S8). MSC-EV administration failed to reduce neurological deficits, infarct volume, brain edema, and neuronal injury in Ly6C^low^ monocyte-deficient mice (Figure 4D-F). Although MSC-EVs increased ICAM-1 expression on ischemic microvessels (Figure 4G), the density of brain-invading CD45^+^ leukocytes, microvascular thrombosis, and BBB permeability remained unchanged (Figure 4H, Figure S9). Peripheral immune analysis (Figure S10A) revealed no significant changes in blood leukocyte composition following MSC-EV treatment in Ly6C^low^ monocyte-deficient mice (Figure S10B), while activated CD4^+^ T cell numbers were increased in the spleen (Figure S10C). Taken together, our data indicate a central role of non-classical Ly6C^low^ monocytes for MSC-EV-induced neuroprotection and immunomodulation after stroke.

**Figure 4.**
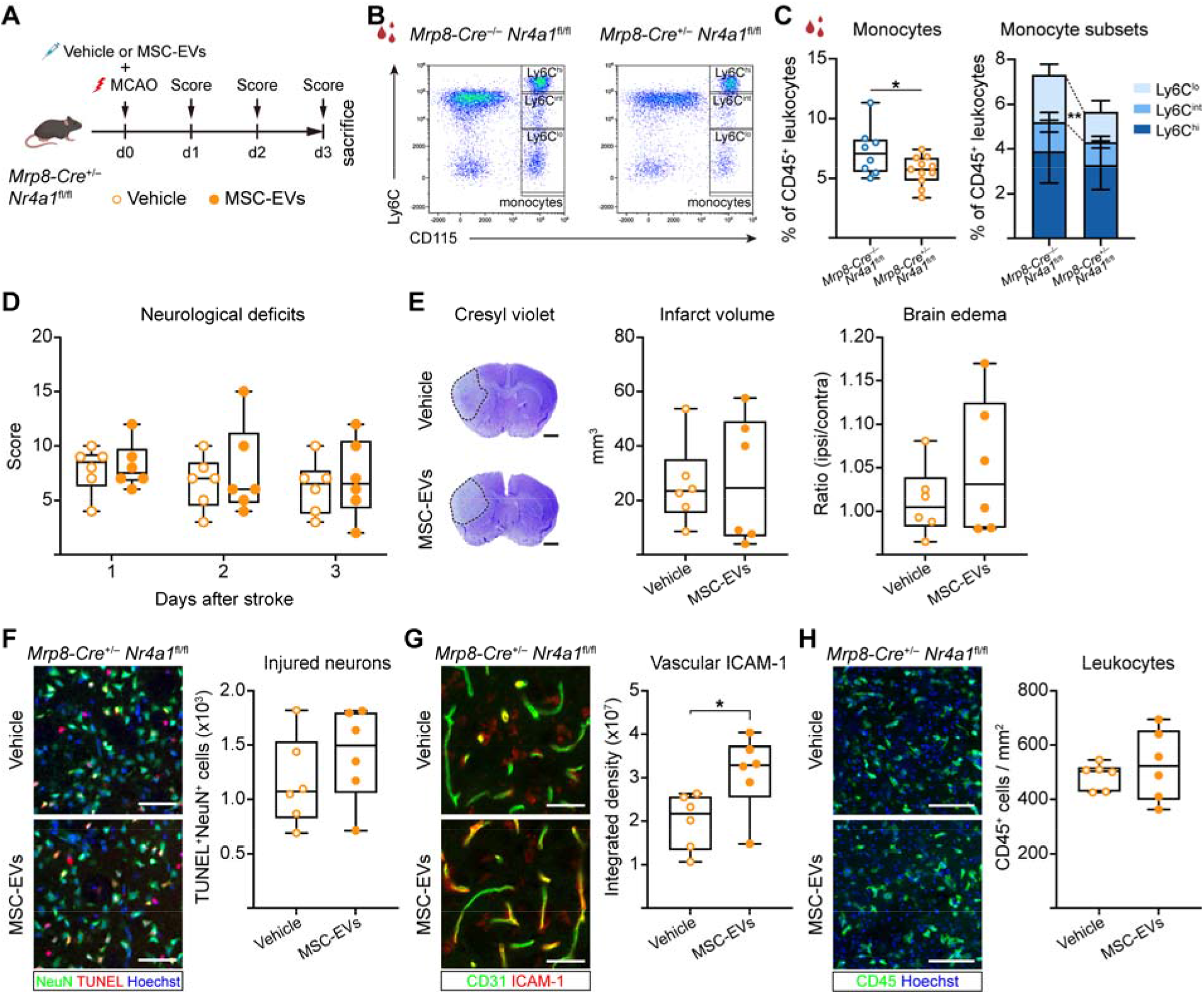
MSC-EVs fail to protect against ischemic injury and reduce brain inflammatory responses in mice deficient in non-classical Ly6C^low^ monocytes. (**A**) Experimental design: Non-classical Ly6C^low^ monocyte deficiency was achieved in *Mrp8-Cre*^+/–^ *Nr4a1*^fl/fl^ mice and compared with *Mrp8-Cre*^−/–^ *Nr4a1*^fl/fl^ mice serving as Ly6C^low^ monocyte competent controls. Following MCAO, vehicle or MSC-EVs (2x10^6^ cell equivalents) were intravenously administered immediately after reperfusion, followed by animal sacrifice at 72 hours post-MCAO (created with https://BioRender.com). (**B**) Gating strategy for circulating monocytes (CD11b^+^ CD115^+^) and (**C**) quantification of monocytes and monocyte subsets in the blood of non-ischemic *Mrp8-Cre*^−/–^ *Nr4a1*^fl/fl^ control and *Mrp8-Cre*^+/–^ *Nr4a1*^*fl*/fl^ mice. (**D**) Neurological deficits evaluated using the Clark score at 24, 48, and 72 hours post-MCAO. (**E**) Infarct volume and brain edema evaluated by cresyl violet staining, (**F**) neuronal injury in the ischemic striatum assessed by TUNEL/NeuN immunolabeling, (**G**) ICAM-1 expression on CD31^+^ microvessels in the ischemic striatum and (**H**) density of brain-infiltrating CD45^+^ leukocytes in the ischemic striatum at 72 hours post-MCAO. Representative cresyl violet stainings (in (**E**)) and immunohistochemistry images (in (**F**-**H**)) are shown. Data were compared by unpaired t test. *p<0.05 (n=6 mice/group). Scale bars: 1 mm (in (**E**)), 50 μm (in (**F, G**)), 100 μm (in (**H**)).

### MSC-EV-induced exacerbation of ischemic injury is mediated by neutrophils and T cells

Since MSC-EV administration induced neutrophil and T cell expansion and overactivation in monocyte-depleted mice, resulting in the exacerbation of ischemic brain damage, we next investigated whether overactivated neutrophils and/ or T cells mediated the exacerbated stroke outcome. In addition to monocyte depletion with clodronate liposomes, we therefore depleted neutrophils or T cells using anti-Ly6G or anti-CD4/CD8 antibody, respectively (Figure 5A). In mice with combined monocyte/ neutrophil and combined monocyte/ T cell depletion, MSC-EV treatment did not worsen neurological deficits, infarct volume, brain edema, neuronal injury, and ICAM-1 abundance (Figure 5B-E), and did not increase brain-infiltrating leukocyte, including neutrophil or T cell counts (Figure 5F, G). Again, microvascular thrombosis and BBB permeability were not recognizably influenced by MSC-EVs (Figure S11). These findings imply a central role of neutrophil and T cell overactivation in the exacerbated stroke outcome induced by MSC-EVs in monocyte-deficient mice.

**Figure 5.**
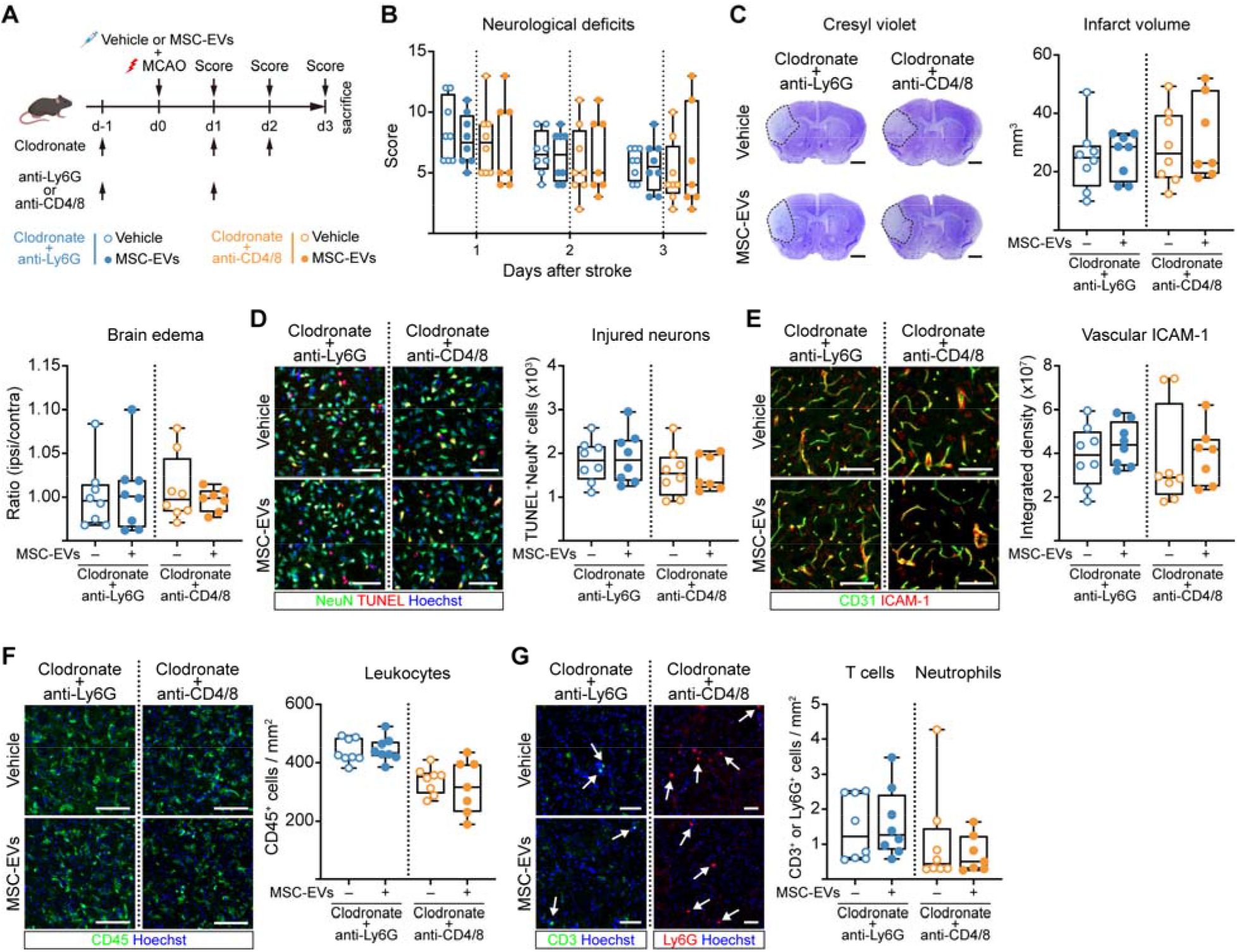
MSC-EV-induced exacerbation of stroke outcome in monocyte-depleted mice is abolished by additional neutrophil or T cell depletion. (**A**) Experimental design: In mice with clodronate liposome-induced monocyte depletion (as above, see Figure 1A), polymorphonuclear neutrophils or T cells were depleted by anti-Ly6G or anti-CD4/CD8 antibody administration 24 hours before and 24 hours after MCAO. Vehicle or MSC-EVs were applied immediately after reperfusion (created with https://BioRender.com). (**B**) Neurological deficits assessed by the Clark score at 24, 48, and 72 hours post-MCAO. (**C**) Infarct volume and brain edema evaluated by cresyl violet staining, (**D**) neuronal injury in the ischemic striatum assessed by TUNEL/NeuN immunolabeling, (**E**) ICAM-1 expression on CD31^+^ microvessels in the ischemic striatum, (**F**) density of brain-infiltrating CD45^+^ leukocytes in the ischemic striatum, and (**G**) density of brain-infiltrating T cells or neutrophils in the ischemic striatum at 72 hours post-MCAO. Representative cresyl violet stainings (in (**C**)) and immunohistochemistry images (in (**D**-**G**)) are shown. Data were compared by unpaired t test or Mann-Whitney U test. No significant group differences were noted (n=7-8 mice/group). Scale bars: 1 mm (in (**C**)), 50 μm (in (**D, G**)), 100 μm (in (**E, F**)).

### MSC-EVs induce functional reprogramming of monocytes in PBMCs from acute ischemic stroke patients

To validate the role of monocytes in mediating MSC-EV-induced immunomodulation across species, we next investigated the effects of MSC-EVs on PBMCs from patients in the acute phase of ischemic stroke. Peripheral blood samples were collected 3–6 days following stroke onset in the prospective NOFF-S cohort.^30^ Intact or monocyte-depleted PBMCs were cultured for 3 days in the presence or absence of MSC-EVs, followed by flow cytometry (Figure 6A, B). MSC-EVs significantly increased CD14 and CD54 expression while decreasing CD16 and HLA-DR expression on monocytes (Figure 6C). In line with previous mouse MCAO studies,^19^ MSC-EVs reduced the frequency of non-classical monocytes (Figure S12A, B), while reducing activated monocyte frequency (Figure 6D). In the absence of monocytes, MSC-EVs increased activated CD4^+^ and CD8^+^ T cell frequencies, along with elevated CD54 and HLA-DR expression (Figure 6E, F). Yet, MSC-EVs did not affect the overall frequencies of CD4^+^ or CD8^+^ T cells (Figure S12C, D). Together, these findings indicate that MSC-EVs promote monocyte conversion to a classical M2-like phenotype and that in the absence of monocytes aberrant T cell activation is induced. Hence, monocytes critically shape the immunomodulatory effects of MSC-EVs in human stroke settings.

**Figure 6.**
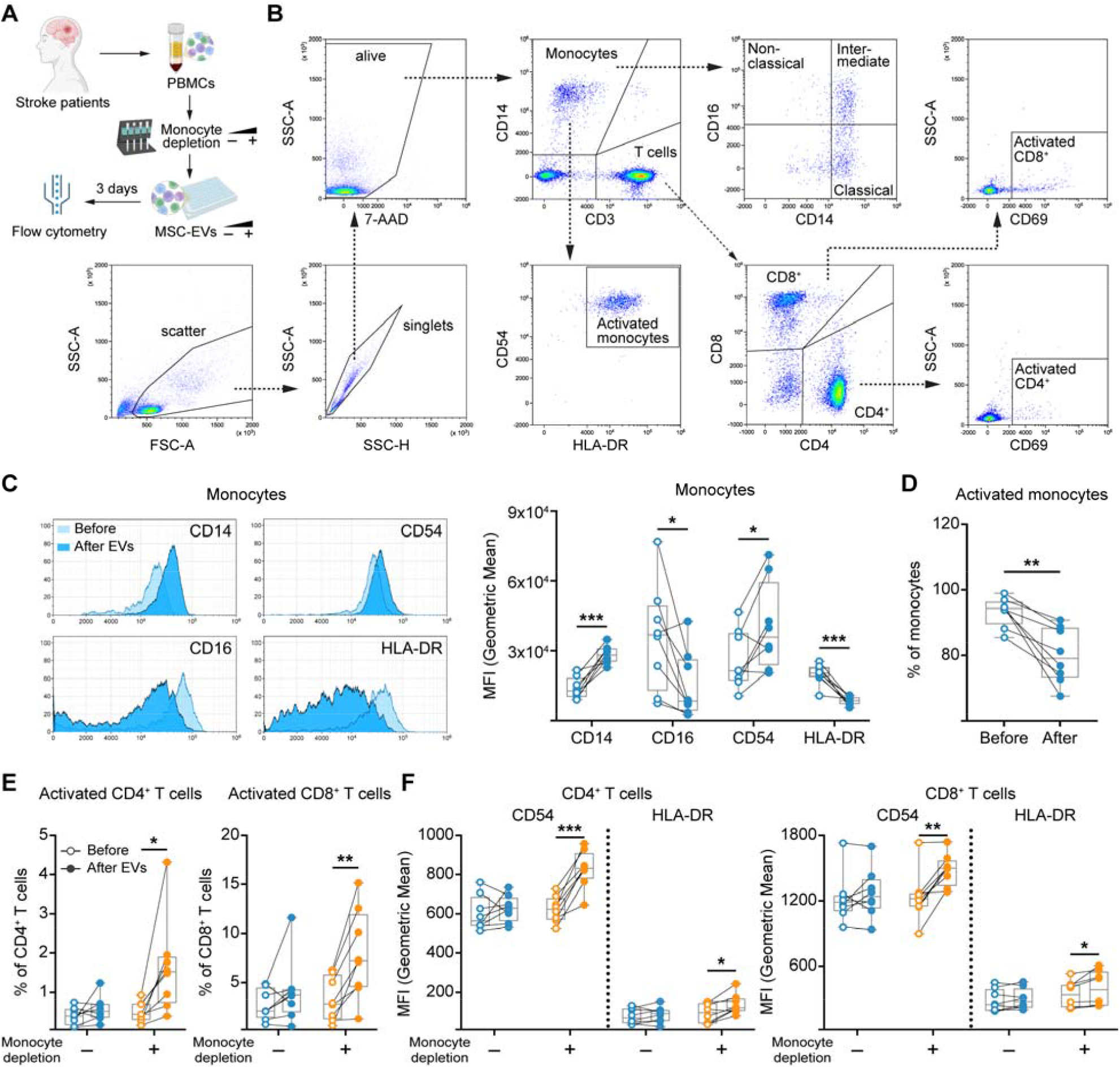
MSC-EVs induce phenotypic monocyte reprogramming while promoting CD4^+^ and CD8^+^ T cell activation in the absence of monocytes in peripheral blood mononuclear cells (PBMCs) obtained from acute ischemic stroke patients. (**A**) Experimental design: PBMCs were isolated from stroke patients 3–6 days after stroke onset. Intact or monocyte-depleted PBMCs were cultured for 3 days in the presence or absence of MSC-EVs, followed by flow cytometric analysis (created with https://BioRender.com). (**B**) Gating strategy for the analysis of PBMCs. (**C**) Representative histograms and corresponding quantification of CD14, CD16, CD54, and HLA-DR expression on monocytes. (**D**) Quantification of activated monocytes (CD14^+^ CD54^+^ HLA-DR^+^), (**E**) activated CD4^+^ (CD3^+^ CD4^+^ CD69^+^) and CD8^+^ (CD3^+^ CD8^+^ CD69^+^) T cells, and (**F**) CD54 and HLA-DR expression levels on CD4^+^ and CD8^+^ T cells in the presence or absence of monocytes, with or without MSC-EV treatment. Data were compared using paired t test. *p<0.05, **p<0.01, ***p<0.001 (n=8 patients; each pair represents an individual stroke patient).

## Discussion

We herein demonstrate that blood monocytes play a critical role in mediating the neuroprotective and immunomodulatory effects of MSC-EVs in mice exposed to MCAO, a clinically relevant ischemic stroke model. Using pharmacological, immunological and genetic approaches to delete monocytes we consistently found that monocyte depletion turned the antiinflammatory effects of MSC-EVs into proinflammatory effects, resulting in the aggravation of ischemic brain damage. Specifically, MSC-EVs increased neutrophil and T cell expansion and activation in the ischemic brain and blood of monocyte-depleted mice, which causally contributed to the injury-exacerbating effects of MSC-EVs, as shown in studies with dual monocyte/ neutrophil and monocyte/ T cell depletion. These findings are clinically relevant: MSC-EVs selectively modulated monocyte differentiation and activation toward a M2-like phenotype in assays using PBMCs from acute ischemic stroke patients, but increased T cell activation when monocytes were absent. Throughout this study, clonally expanded MSCs were used as EV source, which were immortalized using a modified hTERT strategy.^20^ While EVs from primary MSCs exhibit considerable heterogeneity in their immunomodulatory properties due to clonal diversity,^7,10^ this heterogeneity is substantially reduced when ciMSCs are used as EV source.^12,20^

In mice with intact monocyte compartment, MSC-EVs inhibited neutrophil and T cell responses in the ischemic brain of MCAO mice. In monocyte-depleted mice, this inhibitory activity of monocytes was removed, resulting in neutrophil and T cell overactivation in the ischemic brain and blood in response to MSC-EV treatment. *In vitro* studies using PBMCs have previously demonstrated immunomodulatory effects of MSC-EVs on monocytes and lymphocytes, including T and B cells.^13,14,31^ Accumulating evidence indicates that MSC-EV-mediated immunomodulation of T cells occurs indirectly via monocytes.^11,32^ When fluorescently labeled MSC-EVs were added to PBMCs, they were preferentially internalized by monocytes rather than lymphocytes.^13,14^ In *ex vivo* studies using PBMCs obtained from ischemic stroke patients we show that MSC-EVs induce functional monocyte reprogramming reflected by increased CD14 and CD54 expression and reduced CD16 and HLA-DR expression, indicative of the promotion of a classical M2-like phenotype that represses T cell activation. It is well established that classical Ly6C^high^ monocytes can adopt reparative roles in ischemic stroke pathologies by promoting M2 macrophage polarization, eliminating necrotic cell debris, and maintaining brain microvasculature stability.^17,33^ That MSC-EVs may induce an antiinflammatory M2-like monocyte phenotype has previously been shown under conditions unrelated to ischemic injury *in vitro*.^32^ In that earlier study, the promotion of M2-like monocytes went in line with the differentiation of CD4^+^ T cells into Tregs.^32^ Our results suggests that MSC-EVs reprogram monocytes toward a regulatory phenotype after stroke that enhances innate immune sensing while limiting adaptive immune responses.

In our study, MSC-EVs did not influence cerebral microvascular inflammatory responses, indicated by the adhesion molecule ICAM-1, in ischemic brain tissue of monocyte-competent mice, but increased ICAM-1 levels on ischemic microvessels of monocyte-deficient mice. Interacting with CD11b/ CD18 complexes on leukocytes, ICAM-1 mediates the brain tissue infiltration of leukocytes, including neutrophils and T cells.^34^ In our study, ICAM-1 levels increased in response to MSC-EVs in all three monocyte depletion settings, including *Mrp8-Cre*^+/–^ *Nr4a1*^fl/fl^ mice which lacked non-classical Ly6C^low^ patrolling monocytes. Non-classical Ly6C^low^ monocytes control immune responses by patrolling along microvessels.^18^ In a microsphere-induced mouse model of cerebral small vessel disease, non-classical Ly6C^low^ monocytes were found to provide neuroprotection and promote microvascular repair in a CX3CR1 dependent way.^17^ Our results complement these findings by demonstrating that non-classical Ly6C^low^ monocytes constrain microvascular inflammatory responses, preventing the brain entry of leukocytes in MCAO mice. Of note, leukocyte expansion and activation were stronger after combined Ly6C^high^ and Ly6C^low^ monocyte depletion (by clodronate liposomes or anti-CCR2 antibody) than after selective Ly6C^low^ patrolling monocyte deficiency (in *Mrp8-Cre*^+/–^ *Nr4a1*^fl/fl^ mice). Ly6C^high^ inflammatory monocytes can reprogram into Ly6C^low^ antiinflammatory monocytes via Nr4a1 transcription factor activation.^18,35,36^ Thus, in Ly6C^low^ monocyte-deficient mice, Ly6C^high^ monocyte populations may have compensated for the loss of Ly6C^low^ monocytes, attenuating its impact on ischemic brain injury. Importantly, MSC-EVs did not enhance stroke outcome in *Mrp8-Cre*^+/–^ *Nr4a1*^fl/fl^ mice. Our results indicate an important role of non-classical Ly6C^low^ monocytes in the immune responses to MSC-EVs in MCAO mice, possibly as a reservoir for non-classical to classical monocyte differentiation. Indeed, MSC-EVs reduced the population of non-classical CD14^dim^ CD16^+^ monocytes in PBMCs of ischemic stroke patients. An overall reduction of non-classical Ly6C^low^ monocytes by MSC-EVs was previously shown by us in the MCAO mouse brain.^19^ Hence, immune responses to MSC-EVs are remarkably conserved between mice and humans, which highlights the translational relevance of our results.

Collectively, our study identifies monocytes as indispensable intermediaries of MSC-EV-mediated immunomodulation repressing neutrophil and T cell activation, thus enabling the neuroprotective activity of MSC-EVs in ischemic stroke settings. In the forefront of clinical MSC-EV trials, this study emphasizes the need of functional monocyte and T cell assays to evaluate the activity of MSC-EV preparations and rule out side effects or complications.^3,37^ This study advocates thoroughly conducted, possibly personalized activity assays accompanying clinical proof-of-concept trials. Such activity assays will lift cell therapies to the next level.

## Acknowledgments

We thank the Imaging Center Essen (IMCES) at the Faculty of Medicine, University of Duisburg-Essen, Germany, for providing imaging support.

## Supplemental Material

Supplemental Methods

Figures S1-S12

Tables S1-S3

References 38-40

