## Supplemental Material for "Monocytes shape the neuroprotective and immunomodulatory effects of mesenchymal stromal cell-derived extracellular vesicles"

### Supplemental Materials and Methods

#### Legal issues, animal housing, randomization and blinding

Experiments were performed with local government approval (State Office for Consumer Protection and Food North Rhine-Westphalia, Recklinghausen) in accordance to EU (Directive 2010/63/EU) and local institutional guidelines for the care and use of laboratory animals and ARRIVE guidelines 2.0 [23] for experimental data reporting. The experimenter performing the animal experiments and histochemical studies was fully blinded at all stages of the study by another researcher preparing the vehicle and MSC-derived small extracellular vesicle (MSC-EV) solutions. These solutions received dummy names (i.e., solution A, B, C, D), which were unblinded after termination of the study. Animals were kept in a regular inverse 12h:12h light/dark cycle in groups of 5 animals/cage and had free access to food and drinking water. Behavioral tests and animal surgeries were always performed in the morning hours throughout the study.

#### Statistical planning

An a priori sample size calculation was done using an online calculator (https://homepage.univie.ac.at/robin.ristl/samplesize.php?test=ttest). Assuming an alpha error of 5% and a beta error (1–statistical power) of 20%, these calculations determined that 9 animals were needed per group for behavioral and histochemical analyses, provided that EVs modified the mean value by 32.5% and that the standard deviation of the data sample was 25% of the mean value (effect size: 1.3). Whenever a single or up to two animals were missing due to animal dropouts, we did not fill them up. Wherever more animals were lost, animals were complemented by new animals.

#### Expansion and characterization of MSCs

Clonally expanded immortalized MSCs (source 41.5, clone 6) were raised and characterized as we previously described.[21,22] Briefly, MSCs were expanded in low glucose Dulbecco's modified Eagle medium (DMEM; Lonza, Cologne, Germany), supplemented with 10% human platelet lysate (produced in house at Institute of Transfusion Medicine),[38] 100 U/ml penicillin-streptomycin-L-glutamine (Thermo Fisher

Scientific, Waltham, MA, U.S.A.) and 5 IU/ml heparin (Heparin-Natrium-25000, Ratiopharm, Ulm, Germany) and passaged at approximately 80% confluency. In passage 3, MSCs were characterized according to International Society of Cell and Gene Therapy (ISCT) standards.[39] Cell surface phenotypes were analyzed by flow cytometry (CytoFLEX, Software CytExpert 2.3; Beckman-Coulter, Krefeld, Germany) measuring surface antigens including CD14, CD31, CD34, CD44, CD45, CD73, CD90, CD105, and anti-HLA-DR. The osteogenic and adipogenic differentiation potentials of MSCs were confirmed in conventional MSC differentiation assays. In the present study, we used the same clonal MSC line as in our recent study.[21] Starting at passage 3, conditioned media were harvested every 48 hours, centrifuged at 2000 g for 15 minutes to remove cell debris, filtered through filters (0.2 µm), and stored at -20°C until usage. Conditioned media were screened regularly for mycoplasma contamination (VenorGeM OneStep, Minerva Biolabs, Berlin, Germany).

**Preparation and characterization of MSC-EV preparations**

EVs were prepared from thawed pooled conditional media according to our standard procedure, i.e., by polyethylene glycol 6000 (PEG) precipitation followed by ultracentrifugation, as previously described.[7,19,21] PEG precipitation combines volume reduction through water molecule binding with low g-force centrifugation step at 1,500 g for 30 minutes. Pelleted EVs were washed in 0.9% NaCl and re-precipitated by ultracentrifugation at 110,000 g for 130 min (Ti45 rotor, k-factor: 133). MSC-EV samples were dissolved in 10 mM Hepes/0.9% NaCl (Thermo Fisher Scientific) at a concentration of $4x10^7$ cell equivalents per mL and stored at -80°C. Obtained MSC-EV preparations were characterized using our standard procedures as previously described,[7,19] according to the recommendation of minimal information for studies of extracellular vesicles 2018 (MISEV2018).[24] Briefly, concentration and size of MSC-EV preparations were measured by nanoparticle tracking analysis (NTA; Particle Metrix, Meerbusch, Germany) and protein concentrations were determined by a standardized bicinchoninic acid (BCA) assay according the manufacturer's protocol (Pierce, Rockford, IL, U.S.A.). The particle concentration, size, protein concentration, and purity are shown in Table S1. By imaging flow cytometry using the AMNIS ImageStreamX Mark II Flow Cytometer

(Luminex, Seattle, WA, U.S.A.), we had previously shown the presence of $CD9^+$, $CD63^+$ and $CD81^+$ vesicles in our MSC-EV preparations.[21]

**Depletion of monocytes, neutrophils, and T cells**

For unselective depletion of peripheral monocytes/macrophages, phosphate-buffered saline (PBS) loaded control liposomes or clodronate liposomes (Liposoma, Amsterdam, Netherlands) were injected intravenously 24 hours before (50 mg/kg) and 24 and 48 hours after (30 mg/kg) middle cerebral artery occlusion (MCAO) according to previously published protocols.[25,26] Monocyte depletion efficacy by clodronate liposomes was verified by flow cytometry of peripheral blood samples (Figure 1B, C). For depletion of $CCR2^+$ monocytes, 20 µg of IgG isotype antibody (rat IgG2b; BE0090; BioXCell, West Lebanon, NH, U.S.A.) or monoclonal anti-CCR2 antibody MC-21 (provided by Matthias Mack) were injected intravenously 12 hours before and 24 and 48 hours after MCAO.[27] Monocyte depletion efficacy by MC-21 was again verified by flow cytometry of peripheral blood samples (Figure 3B, C). For abolishing non-classical $Ly6C^{low}$ monocytes, *Mrp8-Cre*$^{+/–}$ mice on a C57BL/6J background were crossbred with *Nr4a1*$^{fl/fl}$ mice on a C57BL/6J background (*Nr4a1*$^{flox}$ mice were kindly provided by Pierre Chambon, Institut de Genetique et de Biologie Moleculaire et Cellulaire, Illkirch-Graffenstaden, France) generating *Mrp8-Cre*$^{+/–}$ *Nr4a1*$^{fl/fl}$ mice (generated by Jens Minnerup). The frequency of circulating $Ly6C^{low}$ monocytes was significantly reduced in *Mrp8-Cre*$^{+/–}$ *Nr4a1*$^{fl/fl}$ mice (Figure 4B, C). In defined mouse subgroups, neutrophils were depleted by the intraperitoneal injection of 200 µg anti-Ly6G antibody (BE0075-1; BioXCell) 24 hours before and 24 hours after MCAO,[7] or $CD4^+$ and $CD8^+$ T cells were depleted by the intraperitoneal injection of anti-CD4 (BE0119; BioXCell) and anti-CD8 (BE0061; BioXCell) antibody 24 hours before (100 µg each) and 24 hours after (50 µg each) MCAO.

**Focal cerebral ischemia and MSC-EV treatment**

Focal cerebral ischemia was induced by transient intraluminal MCAO as previously described.[7,19] Briefly, 10-12-week-old male C57BL/6J mice (Harlan Laboratories, Darmstadt, Germany) or *Mrp8-Cre*$^{+/–}$ *Nr4a1*$^{fl/fl}$ mice on a C57BL/6J background (provided by Jens Minnerup) were anesthetized with 1.5% isoflurane (30% $O_2$, remainder $N_2O$). For analgesia, animals were treated subcutaneously with buprenorphine (0.1 mg/kg b.w.; Reckitt Benckiser, Slough, U.K.). Rectal temperature was maintained between 36.5 and 37.0 °C using a feedback-controlled heating system (Fluovac; Harvard apparatus, Holliston, MA, U.S.A.). Cerebral blood flow was recorded by laser Doppler flowmetry using a flexible probe (Perimed, Stockholm, Sweden) attached to the animals' skulls above the core of the middle cerebral artery territory. A midline neck incision was made. The left common and external carotid arteries were isolated and ligated, and the internal carotid artery was temporarily clipped. A silicon-coated 7.0 nylon monofilament (Doccol Corporation, Sharon, MA, U.S.A.) was introduced through a small incision into the common carotid artery and advanced to the carotid bifurcation for MCAO. After 30 min, reperfusion was initiated by monofilament removal. Immediately thereafter, 200 µl normal saline (as vehicle) or $2x10^6$ cell equivalents of MSC-EV preparations dissolved in 200 µl normal saline were administered through the animals' tail vein. Wounds were carefully sutured. For anti-inflammation, animals received injections of carprofen (5 mg/kg; i.p.; twice daily; Bayer Vital, Leverkusen, Germany) during the first 3 days poststroke.

**Analysis of neurological deficits poststroke using the Clark score**

Neurological deficits were evaluated at 24, 48 and 72 hours after MCAO using the Clark score, which evaluates general and focal neurological deficits.[7,19] By adding general and focal deficit scores, the total deficit score was formed.

**Analysis of infarct volume and brain edema by cresyl violet staining**

20-µm-thick coronal brain cryostat sections collected at 1 mm intervals across the forebrain were stained with cresyl violet (C5042; Sigma-Aldrich, Taufkirchen, Germany). In all sections, infarct area was determined using the indirect method that corrects for brain swelling by subtracting the area of healthy tissue of the ischemic hemisphere from that of the contralesional hemisphere using Image J software (National Institute of Health, Bethesda, MD, U.S.A.). Infarct volume was determined by integrating infarct areas from all brain sections. Brain edema was measured as ratio of ipsilateral to contralateral hemisphere volume.

**Immunohistochemical analysis of brain injury, immune infiltrates, microvascular thrombosis and blood-brain barrier permeability**

Coronal brain sections obtained from the rostrocaudal level of the bregma (i.e., the core of the middle cerebral artery territory) were immersed in 0.1 M PBS containing 0.3% Triton X-100 (PBS-T) and 10% normal donkey serum (D9663; Sigma-Aldrich). Sections were incubated overnight at 4°C in monoclonal rabbit anti-NeuN (neuronal nuclear antigen; ab177487; Abcam, Cambridge, U.K.), Alexa Fluor-594 conjugated polyclonal donkey anti-IgG (A21203; Thermo Fisher Scientific), polyclonal goat anti-ICAM-1 (intercellular adhesion molecule-1; AF796; R&D, Minneapolis, MN, U.S.A.), monoclonal rat anti-CD31 (550274; BD Biosciences, Heidelberg, Germany), polyclonal rabbit anti-collagen-IV (AB756P; Merck-Millipore, Darmstadt, Germany), monoclonal rat anti-glycoprotein Ibα (GPIbα) (M043-0; Emfret Analytics, Eibelstadt, Germany), monoclonal rat anti-CD45 (05-1416, Merck Millipore), monoclonal rat anti-Ly6G (lymphocyte antigen 6 locus G; 1A8; 127602; Biolegend, San Diego, CA, U.S.A.) or monoclonal rabbit anti-CD3 (ab16669; Abcam) antibodies. After rinsing, sections were incubated for 1 hour at room temperature in secondary antibodies, as appropriate. Nuclei were counterstained with Hoechst 33342 (62249; Thermo Fisher Scientific). In NeuN stainings, DNA-fragmented, that is, irreversibly injured cells were also detected by terminal deoxynucleotidyl transferase dUTP nick end labeling (TUNEL) (12156792910; Roche Diagnostics, Mannheim, Germany) according to the manufacturer’s protocol. Immunofluorescence stainings were evaluated using a Zeiss AxioOberver.Z1 inverted microscope (Carl Zeiss, Jena, Germany). $NeuN^+$ $TUNEL^+$ neurons, $Ly6G^+$ neutrophils, $CD3^+$ T cells, and $GPIb\alpha^+$ microthrombi were evaluated by counting labeled cells in the entire ischemic striatum. $CD45^+$ leukocytes were evaluated by counting labeled cells in three defined regions of interest (ROI; each measuring 325 µm x 325 µm) in the most lateral part (i.e., core) of the ischemic striatum directly adjacent to the external capsule. For the cell numbers determined in the 3 ROI, mean values were formed. IgG and ICAM-1/CD31 stainings were assessed by measuring optical intensity in the entire ischemic striatum. In the IgG analyses, background stainings were determined in the homologous contralateral non-ischemic striatum and subtracted from values determined in the ischemic striatum.

**Flow cytometry of leukocytes in peripheral blood, spleen, and brain samples**
Single cell suspensions for flow cytometry analysis were obtained as previously described.[7,19] Blood samples were taken from the animals' hearts. After erythrocyte cell lysis with lysis buffer (420302; BioLegend, CA, U.S.A) followed by two washing steps with 0.1 M PBS, blood leukocytes were centrifuged at 500 g for 5 min at 4°C and then stained for 30 minutes at 4°C using antibody cocktails listed in Supplemental Table S2. Following transcardial perfusion with ice-cold 0.1 M PBS, spleen and brain samples were collected. Single cell suspensions of spleens were obtained by meshing the spleen through a 70-µm cell strainer (Life Sciences, New York, NY, U.S.A.) and continuously rinsing with 1% 4-(2-hydroxyethyl)-1-piperazineethanesulfonic acid (HEPES)-buffered Roswell Park Memorial Institute (RPMI)-1640 medium (Thermo Fisher Scientific). Samples were centrifuged at 500 g for 5 min at 4°C. After erythrocyte cell lysis with lysis buffer followed by two washing steps with 0.1 M PBS, spleen leukocytes were centrifuged at 500 g for 5 min at 4°C and then stained for 30 minutes at 4°C using antibody cocktails listed in Supplemental Table S2. Brains were dissected into ischemic (left) and contralateral (right) hemisphere. Single cell suspensions of brains were obtained by meshing the ischemic hemisphere through a 70-µm cell strainer and continuously rinsing with HEPES-buffered RPMI-1640 medium. Samples were centrifuged at 500 g for 5 min at room temperature. After supernatant removal, pellets were resuspended in 37% Percoll (GE Healthcare, Uppsala, Sweden) in 0.1 M PBS. Samples were centrifuged at 900 g for 10 min at room temperature. Myelin and Percoll were aspirated and cell pellets washed twice in 0.1 M PBS. Brain leukocytes were labeled for 30 min at 4°C with antibody cocktails listed in Supplemental Table S2. For intracellular staining of regulatory T cells (Tregs), a Transcription Factor Fixation/Permeabilization Buffer Set (424401; BioLegend, CA, U.S.A) was used according to the manufacturer's instructions. Cell suspensions were analyzed using a

CytoFLEX flow cytometer (Beckman-Coulter) and files were evaluated using Kaluza software V2.2 (Beckman-Coulter).

**Human peripheral blood mononuclear cell (PBMC) isolation and functional assay**

Peripheral blood samples were collected from stroke patients 3–6 days after stroke onset which were enrolled in the NOFF-S study (Neutrophils: Origin, Fate & Function Stroke) recruited via the German Research Foundation–funded Collaborative Research Center TRR332 “Neutrophils: origin, fate and function” (https://www.neutrophils.de, subproject C6; https://drks.de/search/en/trial/DRKS00030825).[30] Baseline characteristics of the stroke patients are summarized in Table S3. PBMCs were obtained by conventional Ficoll density gradient centrifugation as previously reported.[40] For monocyte depletion, PBMCs were subjected to magnetic-activated cell sorting (MACS) using CD14 MicroBeads (130-118-906; Miltenyi Biotec, Bergisch Gladbach, Germany) according to the manufacturer’s instructions. A total of $6 \times 10^5$ viable intact or monocyte-depleted PBMCs were seeded per well in 96-well U-bottom plates (SARSTEDT, Nümbrecht, Germany) in a final volume of 200 µl. Cells were cultured in RPMI 1640 medium (Thermo Fisher Scientific) supplemented with 100 U/ml penicillin, 100 µg/ml streptomycin (both Thermo Fisher Scientific), and 10% human serum (produced in-house) for 3 days at 37 °C in a humidified atmosphere containing 5% $CO_2$ in the presence or absence of MSC-EV preparations ($5x10^4$ cell equivalents dissolved in 5 µl). After culturing, cells were harvested and stained with an antibody cocktail containing anti-CD69, anti-CD3, anti-HLA-DR, anti-CD54, anti-CD16, anti-CD14, anti-CD8, and anti-CD4 antibodies (Table S2). Data acquisition was performed using a CytoFLEX flow cytometer (Beckman Coulter) with CytExpert 2.3 software.

**Statistical analysis**

Statistical analysis was performed using GraphPad Prism (version 10.4.1; GraphPad Software, San Diego, California U.S.A.). Normal distribution was assessed in all datasets using the Shapiro-Wilk normality test. Normally distributed data were analyzed by one-way ANOVA followed by Tukey post hoc test (comparisons between ≥3 groups) or by unpaired or paired two-tailed Student’s t test (comparisons between 2 groups).

Non-normally distributed data were evaluated by Kruskal-Wallis test followed by Dunn's multiple comparisons test (comparisons between ≥3 groups) or two-tailed Mann-Whitney U test (comparisons between 2 groups). Data were presented as box plots with median ± interquartile range with minimum and maximum values as whiskers and individual data points as dots. $P<0.05$ was considered statistically significant.

### Supplemental Figures

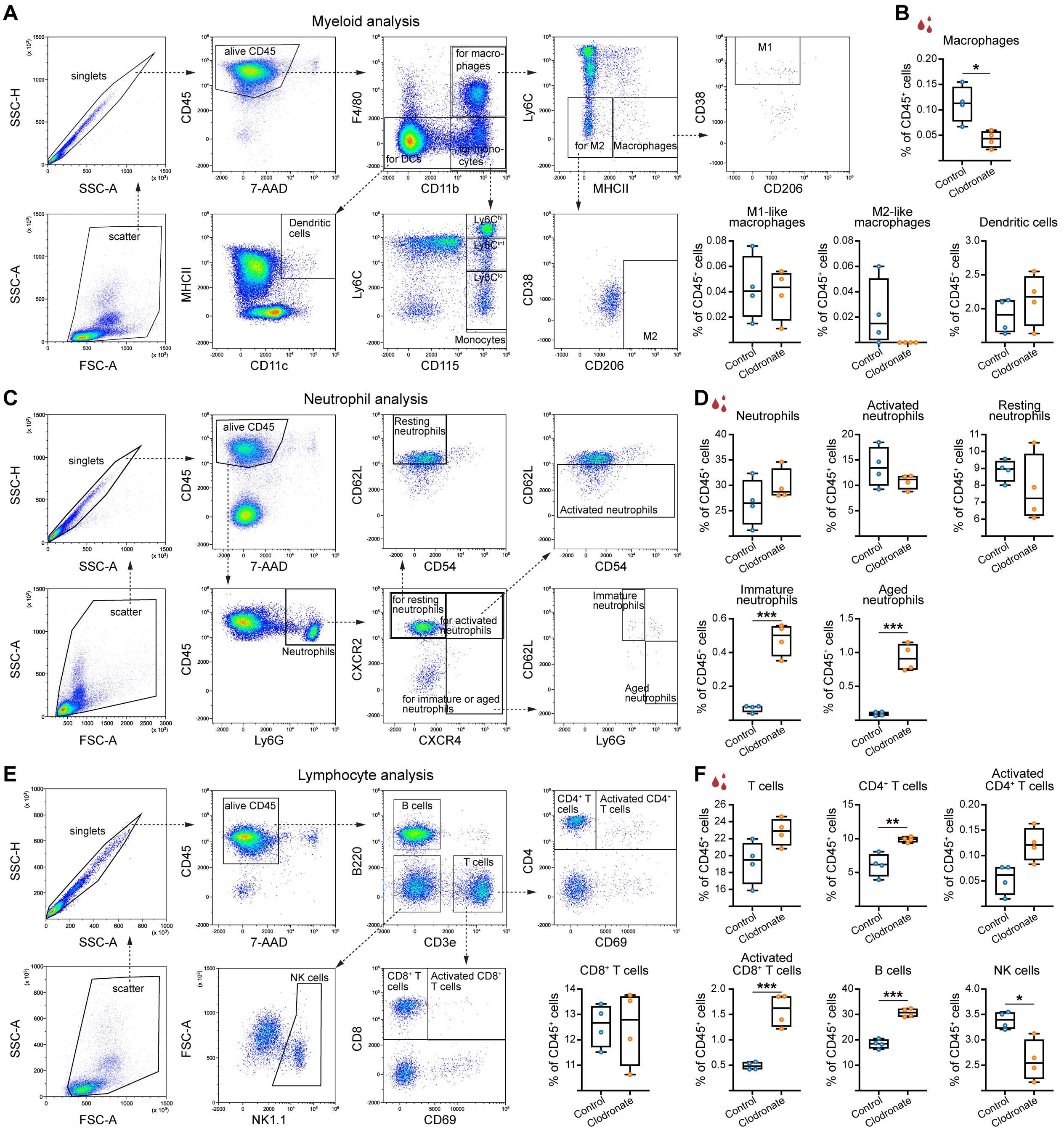


**Figure S1. Expansion and activation of polymorphonuclear neutrophil, T and B cell subsets in response to monocyte depletion by clodronate liposomes in the blood of non-ischemic mice.** Flow cytometry was performed on blood samples from non-ischemic mice 24 hours after intravenous control liposome or clodronate liposome administration. (**A**) Gating strategy for the analysis of myeloid cells within total blood leukocytes. (**B**) Quantification of macrophages (CD11b$^+$ F4/80$^+$ Ly6C$^-$ MHC II$^+$), M1-like macrophages (CD11b$^+$ F4/80$^+$ Ly6C$^-$ MHC II$^+$ CD38$^+$ CD206$^-$), M2-like macrophages (CD11b$^+$ F4/80$^+$ Ly6C$^-$ MHC II$^+$ CD38$^-$ CD206$^+$), and dendritic cells (F4/80$^-$ CD11c$^+$ MHC II$^+$). (**C**) Gating strategy for neutrophil analysis within blood leukocytes. (**D**) Quantification of neutrophils (Ly6G$^+$), activated neutrophils (Ly6G$^+$ CXCR2$^+$ CD62L$^{low}$), resting neutrophils (Ly6G$^+$ CXCR2$^+$ CXCR4$^-$ CD62L$^{high}$ CD54$^{low}$), immature neutrophils (Ly6G$^{low/mid}$ CXCR4$^+$ CD62L$^{high}$), and aged neutrophils (Ly6G$^{high}$ CXCR4$^+$ CD62L$^{low}$). (**E**) Gating strategy for lymphocyte analysis. (**F**) Quantification of T cells (CD3e$^+$), CD4$^+$ T cells (CD3e$^+$ CD4$^+$), activated CD4$^+$ T cells (CD3e$^+$ CD4$^+$ CD69$^+$), CD8$^+$ T cells (CD3e$^+$ CD8$^+$), activated CD8$^+$ T cells (CD3e$^+$ CD8$^+$ CD69$^+$), B cells (CD3e$^-$ B220$^+$), and NK cells (CD3e$^-$ B220$^-$ NK-1.1$^+$). Data were compared by unpaired t test or Mann-Whitney U test. *p<0.05, **p<0.01, ***p<0.001 (n=4 mice/group).

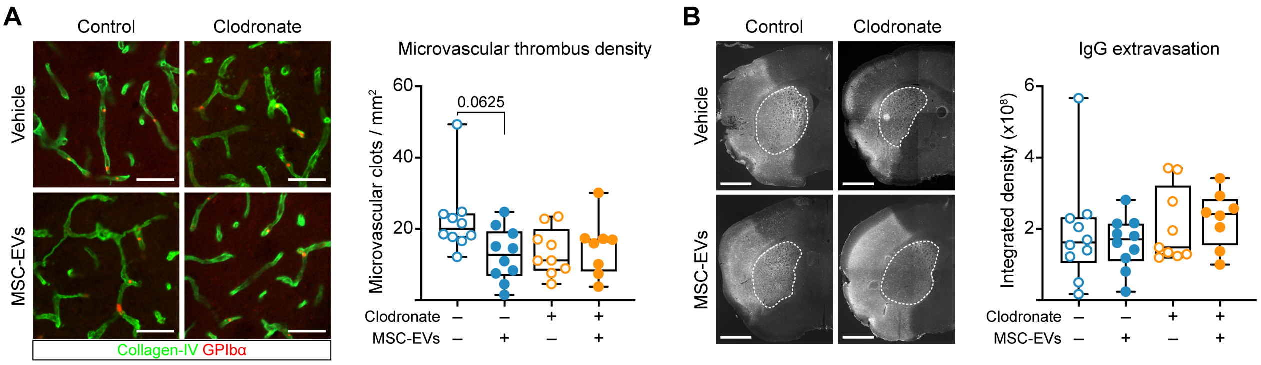


**Figure S2. MSC-EV treatment and monocyte depletion by clodronate liposomes does not influence microvascular thrombosis or blood-brain barrier (BBB) permeability in ischemic brain tissue.** (**A**) Microvascular thrombus density and (**B**) serum IgG extravasation were evaluated by GPIbα/ collagen-IV and IgG immunohistochemistry in the ischemic striatum of mice subjected to transient intraluminal middle cerebral artery occlusion (MCAO). Control or clodronate liposomes were intravenously administered 24 hours prior to MCAO (for details see Figure 1A). Vehicle or MSC-EVs (2x10$^6$ cell equivalents) were intravenously administered immediately after reperfusion. Animals were sacrificed at 72 hours post-MCAO. Representative immunohistochemistry images are shown. Data were compared by Kruskal-Wallis test followed by Dunn’s post hoc test. No significant group differences were noted (n=8-10 mice/group). Scale bars: 50 µm (in (**A**)), 1 mm (in (**B**)).

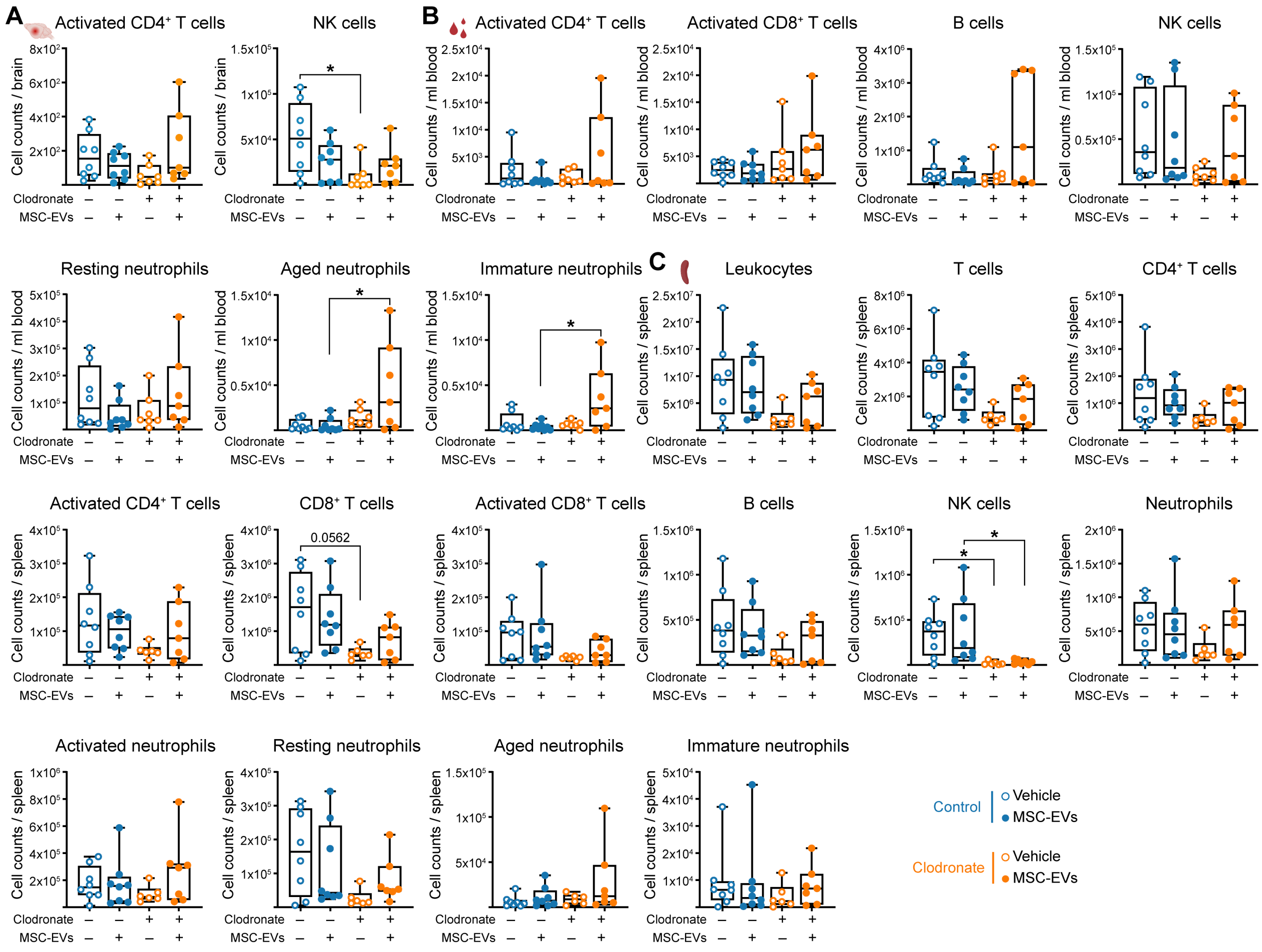

**Figure S3. In the absence of monocytes, MSC-EVs increase aged and immature neutrophils in the blood of ischemic stroke mice.** Monocytes were depleted by control or clodronate liposomes 24 hours prior to MCAO, followed by vehicle or MSC-EV ($2x10^6$ cell equivalents) administration immediately after reperfusion (for details see Figure 1A). Immunophenotyping was performed by flow cytometry on ischemic brain, blood, and spleen samples at 72 hours post-MCAO. (**A**) Quantification of activated $CD4^+$ T cells ($CD3e^+$ $CD4^+$ $CD69^+$) and NK cells ($CD3e^-$ $B220^-$ NK-1.1$^+$) in the ischemic brain. (**B**) Quantification of activated $CD4^+$ T cells ($CD3e^+$ $CD4^+$ $CD69^+$), activated $CD8^+$ T cells ($CD3e^+$ $CD8^+$ $CD69^+$), B cells ($CD3e^-$ $B220^+$), NK cells ($CD3e^-$ $B220^-$ NK-1.1$^+$), resting neutrophils ($Ly6G^+$ $CXCR2^+$ $CXCR4^-$ $CD62L^{high}$ $CD54^{low}$), aged neutrophils ($Ly6G^{high}$ $CXCR4^+$ $CD62L^{low}$), and immature neutrophils ($Ly6G^{low/mid}$ $CXCR4^+$ $CD62L^{high}$) in the blood. (**C**) Quantification of T cells ($CD3e^+$), $CD4^+$ T cells ($CD3e^+$ $CD4^+$), activated $CD4^+$ T cells ($CD3e^+$ $CD4^+$ $CD69^+$), $CD8^+$ T cells ($CD3e^+$ $CD8^+$), activated $CD8^+$ T cells ($CD3e^+$

$CD8^+$ $CD69^+$), B cells ($CD3e^−$ $B220^+$), NK cells ($CD3e^−$ $B220^−$ $NK\text{-}1.1^+$), neutrophils ($Ly6G^+$), activated neutrophils ($Ly6G^+$ $CXCR2^+$ $CD62L^{low}$), resting neutrophils ($Ly6G^+$ $CXCR2^+$ $CXCR4^−$ $CD62L^{high}$ $CD54^{low}$), aged neutrophils ($Ly6G^{high}$ $CXCR4^+$ $CD62L^{low}$), and immature neutrophils ($Ly6G^{low/mid}$ $CXCR4^+$ $CD62L^{high}$) in the spleen. Data were compared by Kruskal-Wallis test followed by Dunn's post hoc test or one-way ANOVA followed by Tukey post hoc test. *$p<0.05$ (n=7-8 mice/group).

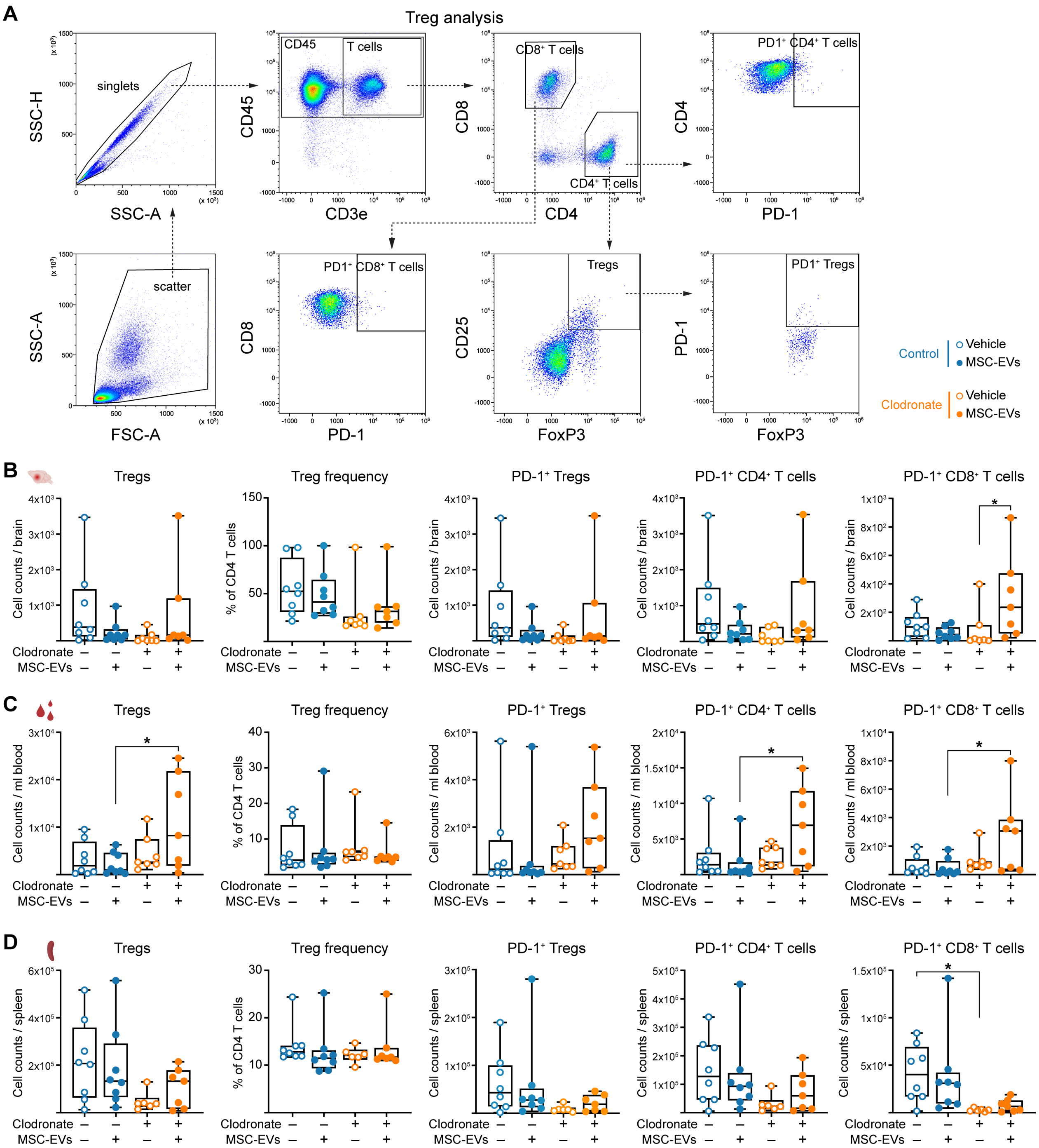


**Figure S4. MSC-EV-induced exacerbation of ischemic injury in monocyte-depleted mice is associated with T cell suppression/ exhaustion phenotypes.** Monocytes were depleted by control or clodronate liposomes 24 hours prior to MCAO, followed by vehicle or MSC-EV (2x10$^6$ cell equivalents) administration immediately after reperfusion (for details see Figure 1A). T cell suppressive/ exhausted phenotypes were analyzed by flow cytometry in ischemic brain, blood, and spleen samples from mice sacrificed at 72 hours post-MCAO. (**A**) Gating strategy for T cell immunosuppression/ exhaustion analysis. Quantification of regulatory T cells (Tregs) ($CD3e^+$ $CD4^+$ $CD25^+$ $FoxP3^+$), PD-$1^+$ Tregs ($CD3e^+$ $CD4^+$ $CD25^+$ $FoxP3^+$ PD-$1^+$), PD-$1^+$ $CD4^+$ T cells ($CD3e^+$ $CD4^+$ PD-$1^+$), PD-$1^+$ $CD8^+$ T cells ($CD3e^+$ $CD8^+$ PD-$1^+$), and Treg frequency (percentage of $CD4^+$ T cells) in (**B**) the ischemic brain, (**C**) the blood, and (**D**) the spleen. Data were compared by Kruskal-Wallis test followed by Dunn’s post hoc test or one-way ANOVA followed by Tukey post hoc test. *$p<0.05$ (n=7-8 mice/group).

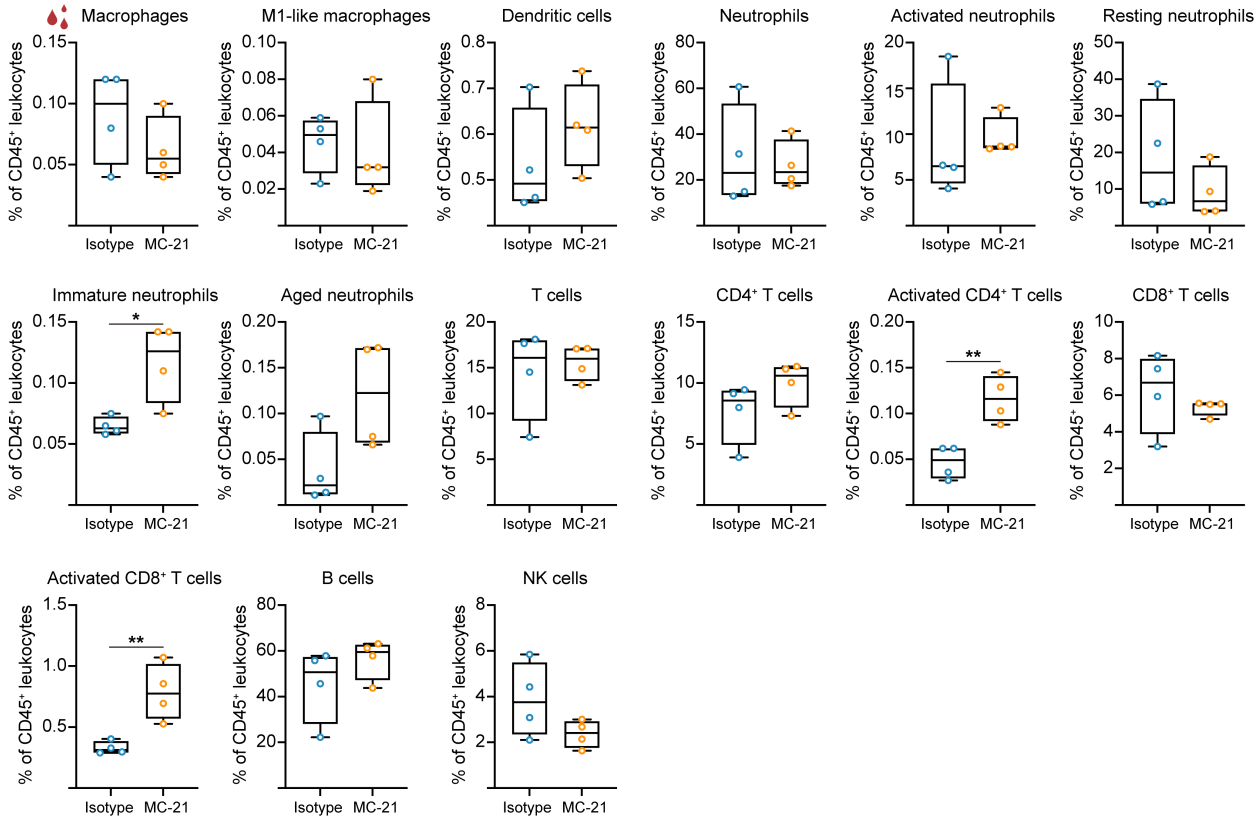


**Figure S5. Monocyte depletion by anti-CCR2 antibody (MC-21) increases immature neutrophils, activated CD4$^{+}$ T cells and activated CD8$^{+}$ T cells in the blood of non-ischemic mice.** Flow cytometric analysis was performed on blood samples of non-ischemic mice 12 hours after intravenous isotype control or MC-21 antibody administration. Blood leukocyte subsets, including macrophages, M1-like macrophages, dendritic cells, neutrophils, activated neutrophils, resting neutrophils, immature neutrophils, aged neutrophils, T cells, CD4$^{+}$ T cells, activated CD4$^{+}$ T cells, CD8$^{+}$ T cells, activated CD8$^{+}$ T cells, B cells, and NK cells, were quantified. Data were compared by unpaired t test or Mann-Whitney U test. *$p<0.05$, **$p<0.01$ (n=4 mice/group).

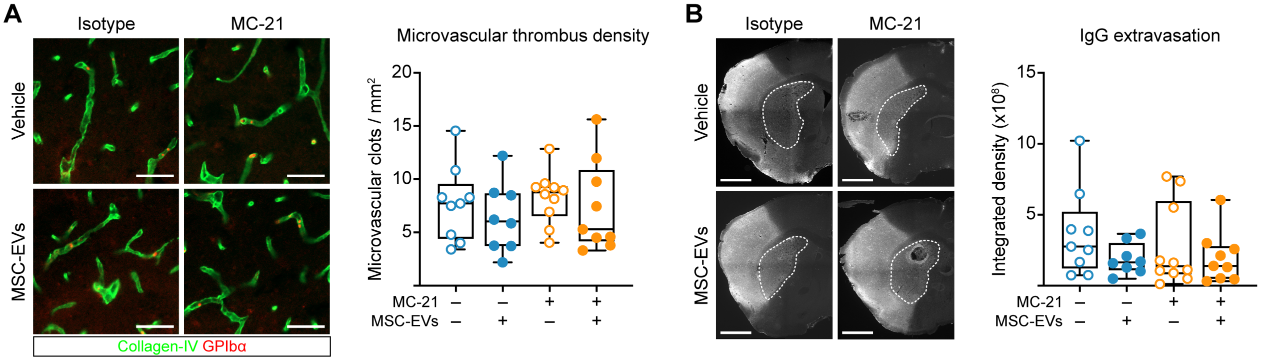


**Figure S6. MSC-EV treatment and monocyte depletion by anti-CCR2 antibody MC-21 does not influence microvascular thrombosis or BBB permeability in ischemic brain tissue.** (**A**) Microvascular thrombus density and (**B**) IgG extravasation were evaluated by immunohistochemistry in the ischemic striatum of mice subjected to MCAO. Isotype control antibody or anti-CCR2 antibody MC-21 were intravenously applied 12 hours prior to MCAO (for details see Figure 3A). Vehicle or MSC-EVs ($2x10^6$ cell equivalents) were intravenously administered immediately after reperfusion. Animals were sacrificed at 72 hours post-MCAO. Representative immunohistochemistry images are shown. Data were compared by Kruskal-Wallis test followed by Dunn's post hoc test. No significant group differences were noted (n=8-10 mice/group). Scale bars: 50 μm (in (**A**)), 1 mm (in (**B**)).

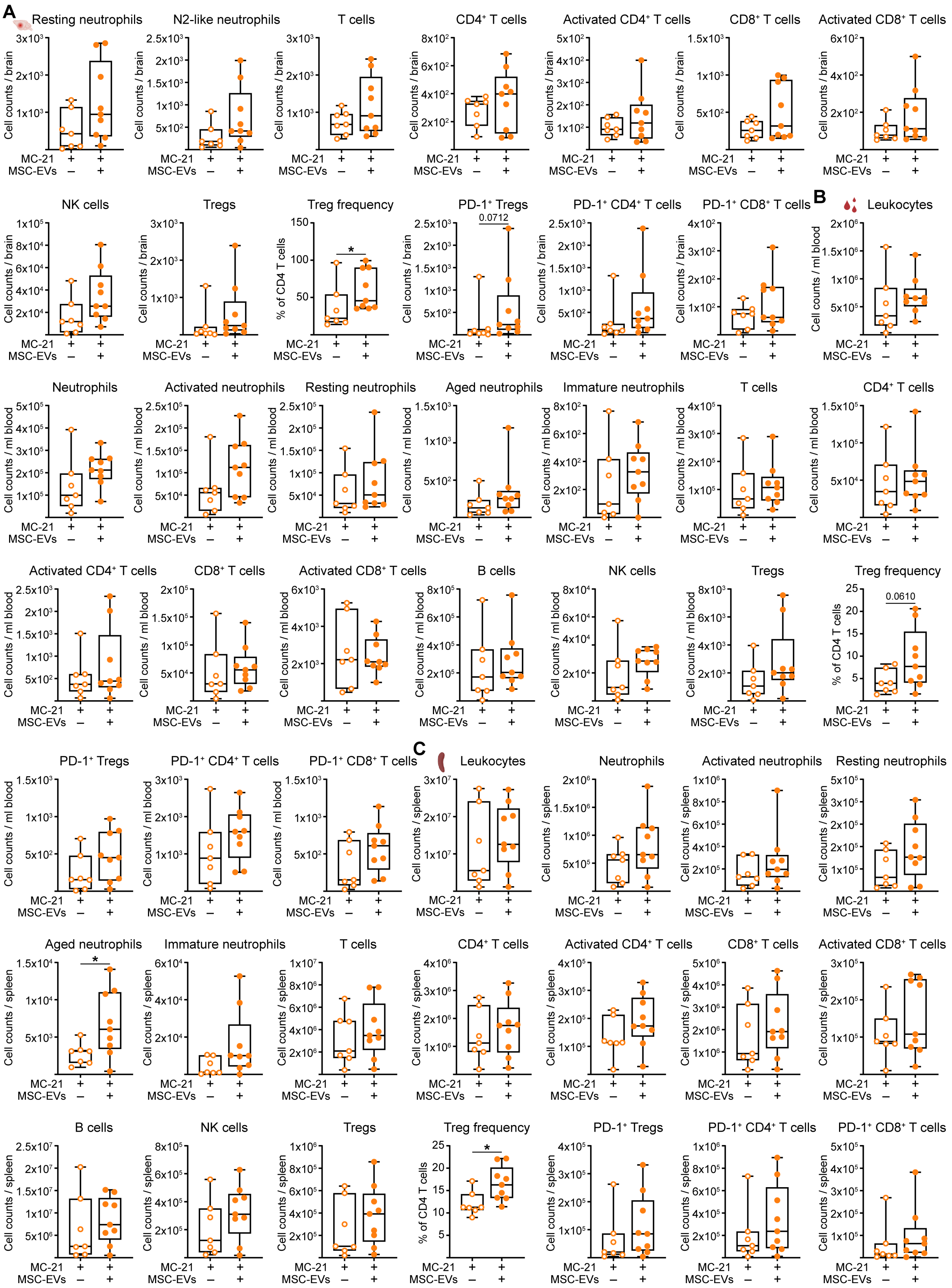
A
Resting neutrophils
N2-like neutrophils
T cells
CD4⁺ T cells
Activated CD4⁺ T cells
CD8⁺ T cells
Activated CD8⁺ T cells
NK cells
Tregs
Treg frequency
PD-1⁺ Tregs
0.0712
PD-1⁺ CD4⁺ T cells
PD-1⁺ CD8⁺ T cells
Cell counts / brain
% of CD4 T cells
*
B
Leukocytes
Neutrophils
Activated neutrophils
Resting neutrophils
Aged neutrophils
Immature neutrophils
T cells
CD4⁺ T cells
Activated CD4⁺ T cells
CD8⁺ T cells
Activated CD8⁺ T cells
B cells
NK cells
Tregs
Treg frequency
0.0610
PD-1⁺ Tregs
PD-1⁺ CD4⁺ T cells
PD-1⁺ CD8⁺ T cells
Cell counts / ml blood
C
Leukocytes
Neutrophils
Activated neutrophils
Resting neutrophils
Aged neutrophils
Immature neutrophils
T cells
CD4⁺ T cells
Activated CD4⁺ T cells
CD8⁺ T cells
Activated CD8⁺ T cells
B cells
NK cells
Tregs
Treg frequency
PD-1⁺ Tregs
PD-1⁺ CD4⁺ T cells
PD-1⁺ CD8⁺ T cells
Cell counts / spleen
MC-21 + +
MSC-EVs – +

**Figure S7. MSC-EV-induced exacerbation of ischemic brain injury in $CCR2^+$ monocyte-depleted mice is associated with increased Treg frequency, suggestive of a compensatory negative feedback mechanism.** Anti-CCR2 antibody MC-21 were injected 12 hours prior to MCAO for monocyte depletion (for details see Figure 3A). Vehicle or MSC-EVs ($2x10^6$ cell equivalents) were intravenously delivered immediately after reperfusion. Flow cytometry was performed on ischemic brain, blood, and spleen samples 72 hours post-MCAO. Leukocyte subsets and Treg frequency were quantified in (**A**) the ischemic brain, (**B**) the blood, and (**C**) the spleen. Data were compared by unpaired t test or Mann-Whitney U test. $*p<0.05$ (n=7-9 mice/group).

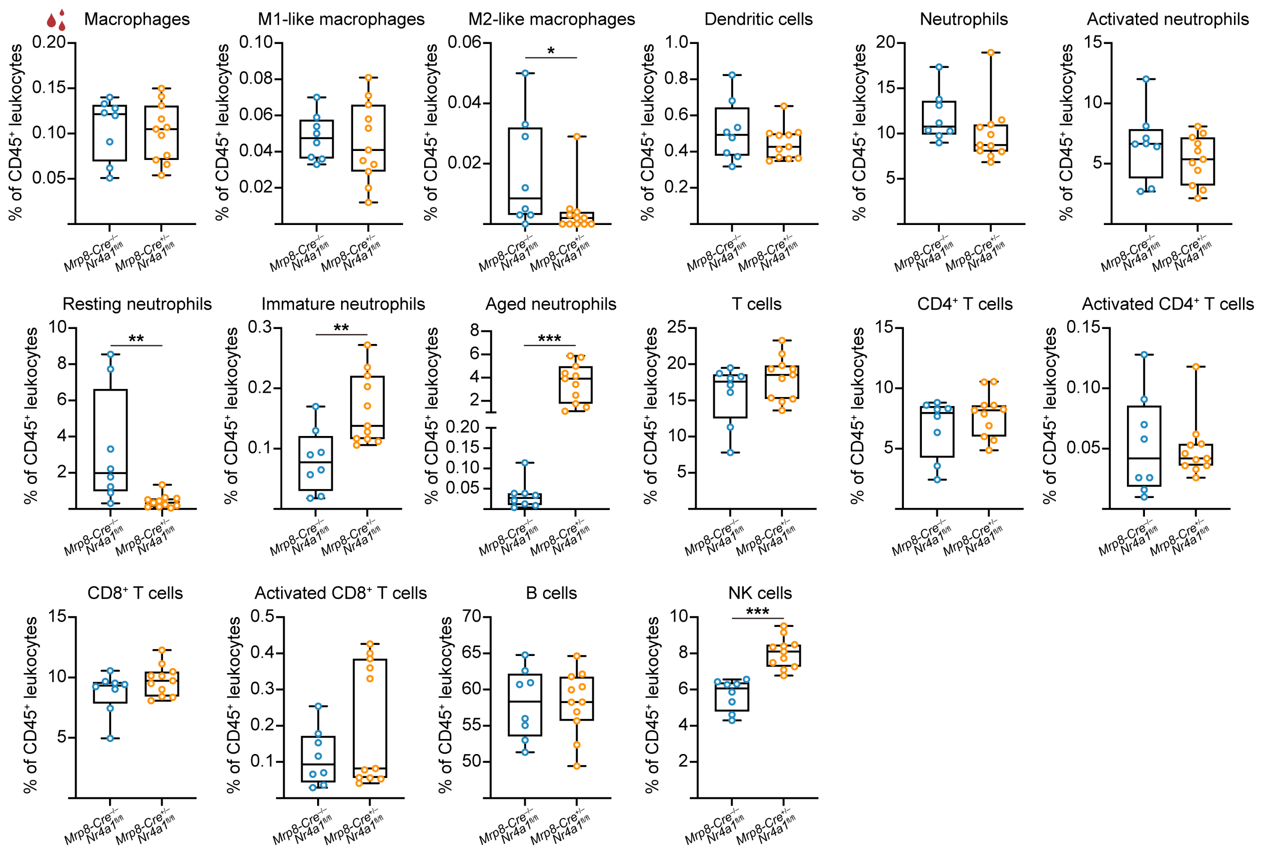


**Figure S8. Immune phenotype of blood leukocytes other than monocytes of Ly6C$^{low}$ deficient *Mrp8-Cre*$^{+/–}$ *Nr4a1*$^{fl/fl}$ mice.** Flow cytometric quantification of leukocyte subsets, namely macrophages, M1-like macrophages, M2-like macrophages, dendritic cells, neutrophils, activated neutrophils, resting neutrophils, immature neutrophils, aged neutrophils, T cells, CD4$^{+}$ T cells, activated CD4$^{+}$ T cells, CD8$^{+}$ T cells, activated CD8$^{+}$ T cells, B cells, and NK cells in the blood of non-ischemic *Mrp8-Cre*$^{–/–}$ *Nr4a1*$^{fl/fl}$ control and *Mrp8-Cre*$^{+/–}$ *Nr4a1*$^{fl/fl}$ mice. Data were compared by unpaired t test or Mann-Whitney U test. *p<0.05, **p<0.01, ***p<0.001 (n=8-11 mice/group).

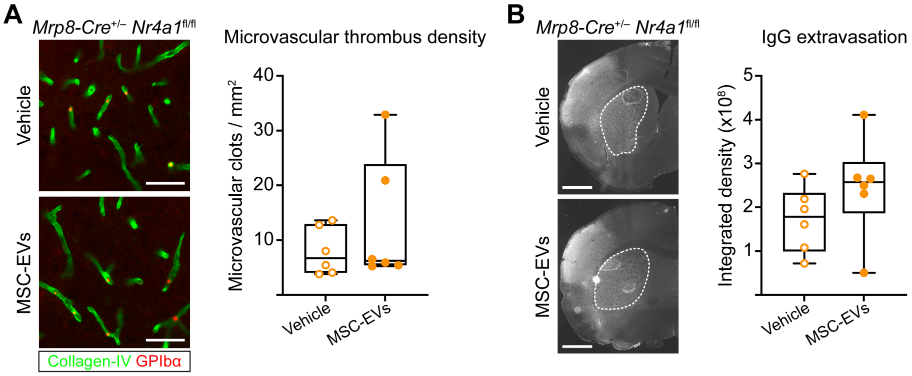


**Figure S9. MSC-EV treatment does not affect microvascular thrombosis or BBB permeability in ischemic brain tissue of *Mrp8-Cre*$^{+/-}$ *Nr4a1*$^{fl/fl}$ mice deficient for non-classical Ly6C$^{low}$ monocytes.** (**A**) Microvascular thrombus density and (**B**) serum IgG extravasation were evaluated by immunohistochemistry in the ischemic striatum of *Mrp8-Cre*$^{+/-}$ *Nr4a1*$^{fl/fl}$ mice exposed to MCAO. Vehicle or MSC-EVs (2x10$^6$ cell equivalents) were intravenously administered immediately after reperfusion. Animals were sacrificed at 72 hours post-MCAO. Representative immunohistochemistry images are shown. Data were compared by Mann-Whitney U test (in (**A**)) or unpaired t test (in (**B**)). No significant group differences were noted (n=6 mice/group). Scale bars: 50 µm (in (**A**)), 1 mm (in (**B**)).

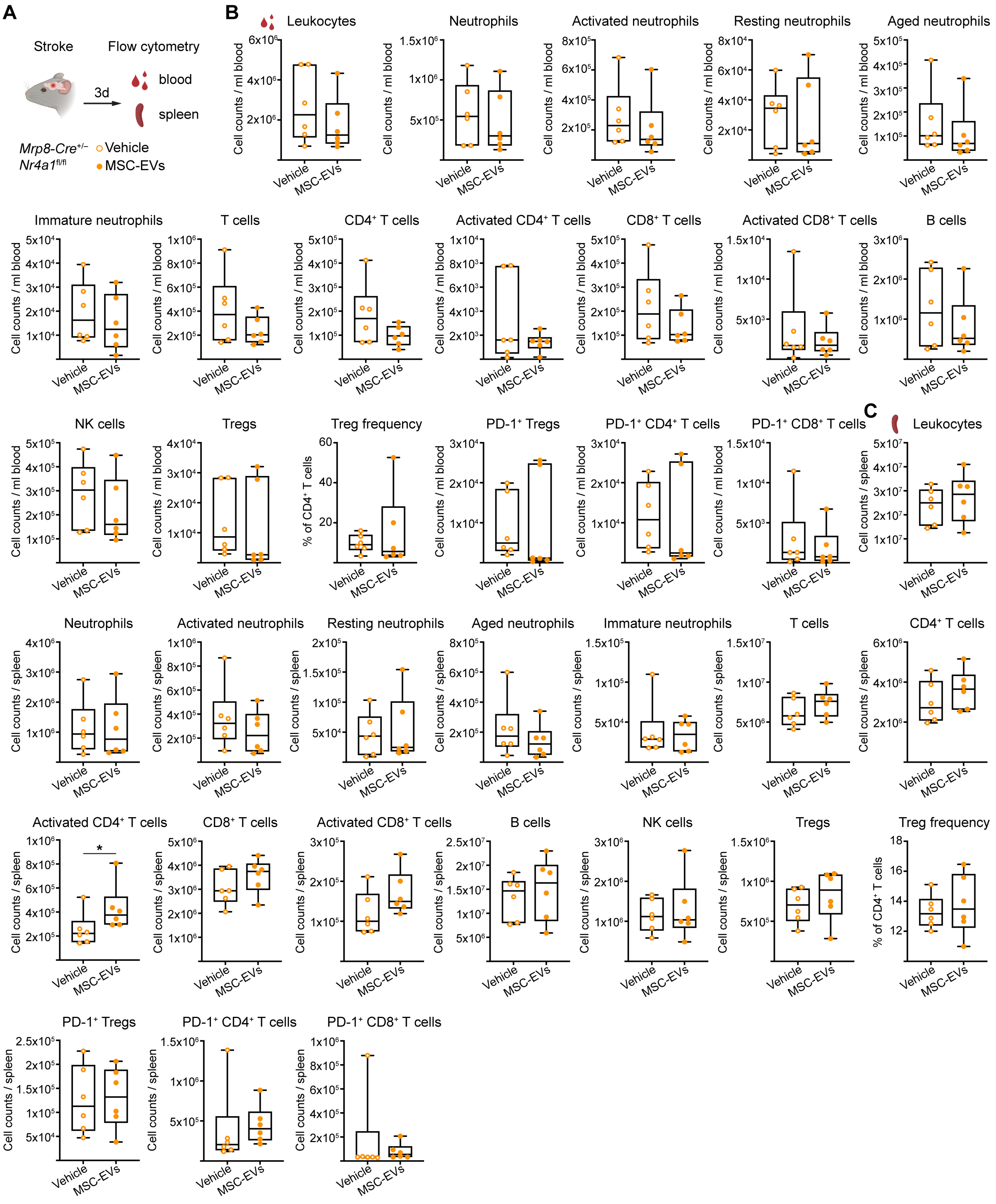


**Figure S10. CD4⁺ T cells are overactivated by MSC-EVs in the spleen of ischemic *Mrp8-Cre*$^{+/–}$ *Nr4a1*$^{fl/fl}$ mice deficient for non-classical Ly6C$^{low}$ monocytes.** (A)

Experimental design: Flow cytometry of blood and spleen samples of *Mrp8-Cre*$^{+/-}$ *Nr4a1*$^{fl/fl}$ mice exposed to MCAO. Vehicle or MSC-EVs ($2x10^6$ cell equivalents) were intravenously administered immediately after reperfusion. Animals were sacrificed at 72 hours post-MCAO (created with https://BioRender.com). Quantification of leukocyte subsets in (**B**) the blood and (**C**) the spleen. Data were compared by unpaired t test or Mann-Whitney U test. *p<0.05 (n=6 mice/group).

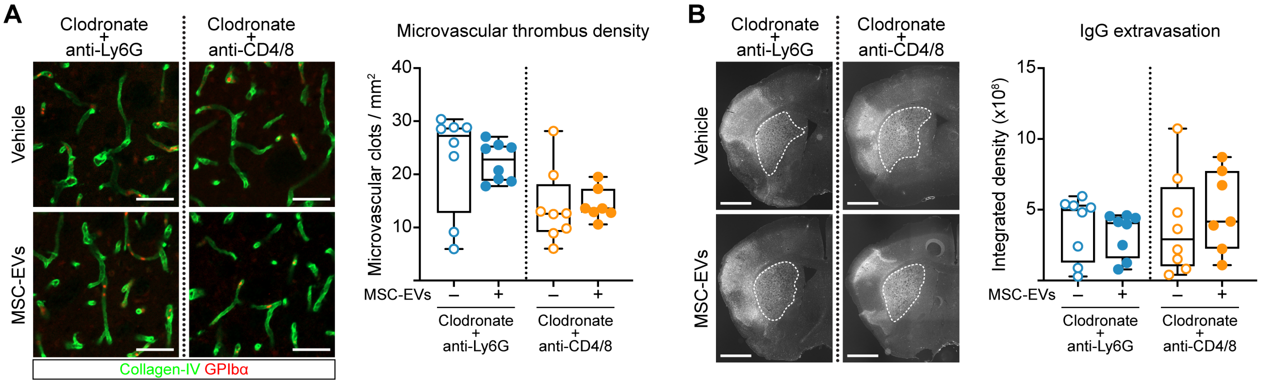


**Figure S11. MSC-EV treatment does not influence microvascular thrombosis or BBB permeability in ischemic brain tissue of mice with combined monocyte and neutrophil depletion or monocyte and T cell depletion.** (**A**) Microvascular thrombus density and (**B**) serum IgG extravasation were evaluated by immunohistochemistry in the ischemic striatum of transient MCAO mice. Control or clodronate liposomes were intravenously administered 24 hours prior to MCAO for monocyte depletion. Anti-Ly6G or anti-CD4/CD8 antibody were applied for neutrophil or T cell depletion 24 hours prior to MCAO (for details see Figure 5A). Vehicle or MSC-EVs (2x10$^6$ cell equivalents) were intravenously delivered immediately after reperfusion. Animals were sacrificed at 72 hours post-MCAO. Representative immunohistochemistry images are shown. Data were compared by Mann-Whitney U test (in (**A**)) or unpaired t test (in (**B**)). No significant group differences were noted (n=7-8 mice/group). Scale bars: 50 µm (in (**A**)), 1 mm (in (**B**)).

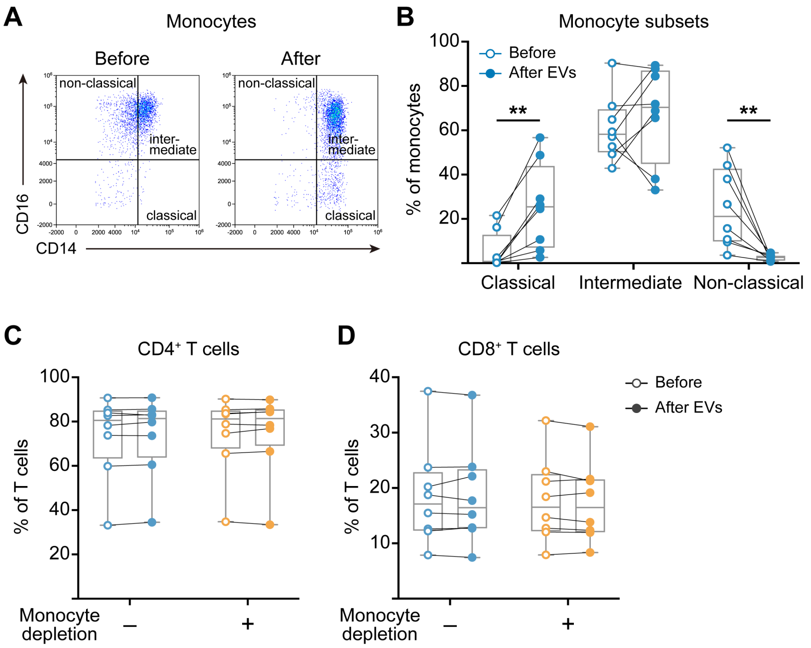


**Figure S12. MSC-EVs reprogram monocytes toward a classical-like phenotype while do not influence CD4$^+$ and CD8$^+$ T cell frequencies in peripheral blood mononuclear cells (PBMCs) from patients with acute ischemic stroke.** (**A**) Gating strategy for monocytes in PBMCs classified as classical (CD14$^{high}$ CD16$^-$), intermediate (CD14$^{high}$ CD16$^+$), and non-classical (CD14$^{dim}$ CD16$^+$) monocytes. Flow cytometric quantification of (**B**) monocyte subsets, (**C**) CD4$^+$ T cells and (**D**) CD8$^+$ T cells in intact or monocyte-depleted PBMCs after 3-day culture with or without MSC-EVs. Data were compared by paired t test. **p<0.01 (n=8 patients; each pair represents an individual patient).

### Supplementary Tables

#### Supplemental Table S1: Biophysical characteristics of the applied MSC-EV preparations.

| | Particle concentration [particles/ ml] | Particle size [nm] | Protein concentration [µg/ µl] | Purity [particles/ mg protein] |
|---|---|---|---|---|
| MSC-EVs (source 41.5) | $1.9 \times 10^{11}$ | 109.6 | 4.7 | $4.04 \times 10^{10}$ |

#### Supplemental Table S2: Antibodies used for flow cytometry.

| Antigen | Conjugate | Host/isotype | Clone | Supplier |
|---|---|---|---|---|
| Mouse CD45 | Pacific blue | Rat IgG2b, kappa | 30F11 | BioLegend |
| Mouse CD45 | BV605 | Rat IgG2b, kappa | 30F11 | BioLegend |
| Mouse Ly6G | PE | Rat IgG2a, kappa | 1A8 | BioLegend |
| Mouse CXCR2 | BV786 | Rat IgG2b, kappa | V48-2310 | BD Biosciences |
| Mouse CXCR4 | FITC | Recombinant human IgG1 | REA107 | Miltenyi Biotec |
| Mouse PD-L1 | PE/Dazzle 594 | Rat IgG2a, kappa | 10F.9G2 | BioLegend |
| Mouse CD170 | APC | Rat IgG2a, kappa | S17007L | BioLegend |
| Mouse CD54 (ICAM-1) | APC-Vio 770 | Recombinant human IgG1 | REA171 | Miltenyi Biotec |
| Mouse CD62L | eFluor 450 | Rat IgG2a, kappa | MEL-14 | Thermo Fisher Scientific |
| Mouse Ly6C | FITC | Rat IgM, kappa | AL21 | BD Biosciences |
| Mouse CD11c | PE/Dazzle 594 | Hamster IgG | N418 | BioLegend |
| Mouse CD115 | PE-Cy7 | Rat IgG2a, kappa | AFS98 | Thermo Fisher Scientific |
| Mouse CD11b | APC | Rat IgG2b, kappa | M1/70 | Thermo Fisher Scientific |
| Mouse F4/80 | APC/Cy7 | Rat IgG2a, kappa | BM8 | BioLegend |
| Mouse CD38 | BV421 | Rat IgG2a, kappa | Ab90 | BD Biosciences |
| Mouse MHCII | VioGreen | Recombinant human IgG1 | REA813 | Miltenyi Biotec |
| Mouse CD206 | BV785 | Rat IgG2a, kappa | C068C2 | BioLegend |

| | | | | |
|---|---|---|---|---|
| Mouse CD3ε | Alexa Fluor 647 | Hamster IgG | 145-2C11 | BioLegend |
| Mouse CD4 | BV605 | Rat IgG2a, kappa | RM4-5 | BD Biosciences |
| Mouse CD8α | BV786 | Rat IgG2a, kappa | 53-6.7 | BD Biosciences |
| Mouse B220 | PE | Rat IgG2a, kappa | RA3-6B2 | BD Biosciences |
| Mouse NK-1.1 | FITC | Rat IgG2a, kappa | PK136 | BD Biosciences |
| Mouse CD69 | PE-Cy7 | Hamster IgG | H1.2F3 | BioLegend |
| Mouse CD25 | FITC | Rat IgM, kappa | 7D4 | Miltenyi Biotec |
| Mouse PD-1 | PerCP-Vio 770 | Recombinant human IgG1 | REA802 | Miltenyi Biotec |
| Mouse Foxp3 | PE | Rat IgG2a, kappa | FJK-16s | Thermo Fisher Scientific |
| Human CD69 | FITC | Mouse IgG1, kappa | FN50 | BioLegend |
| Human HLA-DR | ECD | Mouse IgG1 | Immu-357 | Beckman Coulter |
| Human CD3 | APC | Mouse BALB/c IgG1, kappa | SK-7 | BD Biosciences |
| Human CD54 | Alexa Fluor 700 | Mouse IgG2b | 1H4 | Exbio |
| Human CD16 | APC-eFluo 780 | Mouse IgG1, kappa | eBioCB16 (CB16) | Invitrogen |
| Human CD14 | Pacific Orange | Mouse IgG1, kappa | MEM-15 | Exbio |
| Human CD8 | BV650 | Mouse IgG1, kappa | SK1 | BioLegend |
| Human CD4 | BV785 | Mouse IgG1, kappa | RPA-T4 | BioLegend |

**Supplemental Table S3: Baseline characteristics for ischemic stroke patients from the NOFF-S study.**

| | Total cohort (n=8 patients) |
|---|---|
| Age | 70.5 ± 15.3 |
| Female sex | 3 (37.5) |
| NIHSS score at admission, median [Q1, Q3] | 2.0 [1.0, 6.0] |
| **Stroke subtype** | |
| Large artery atherosclerosis | 3 (37.5) |

| | |
|---|---|
| Cardioembolic | 0 (0.0) |
| Small artery occlusion | 1 (12.5) |
| Other causes | 0 (0.0) |
| Undefined | 4 (50.0) |
| **Cardiovascular risk factors** | |
| Arterial hypertension | 7 (87.5) |
| Diabetes mellitus | 3 (37.5) |
| Dyslipidemia | 3 (37.5) |
| Atrial fibrillation | 0 (0.0) |
| Current smoking | 2 (25.0) |
| BMI | 28.3 ± 9.1 |
| Regular alcohol consumption | 4 (50.0) |
| Physical inactivity | 6 (75.0) |
| **Cardiovascular diseases** | |
| History of myocardial infarction | 1 (12.5) |
| Coronary artery disease | 2 (25.0) |
| Peripheral artery disease | 1 (12.5) |
| Heart failure | 1 (12.5) |
| History of transient ischemic attack | 0 (0.0) |
| Carotid dissection | 1 (12.5) |
| Internal carotid artery stenosis >50% | 0 (0.0) |
| **Intervention** | |
| Thrombolysis | 2 (25.0) |
| Thrombectomy | 0 (0.0) |
| Thrombolysis plus thrombectomy | 0 (0.0) |
| None | 6 (75.0) |
| **Cardiovascular medication** | |
| Antihypertensive drugs | 6 (75.0) |

| | |
|---|---|
| Antidiabetics | 2 (25.0) |
| Lipid-modifying agents | 4 (50.0) |
| Antithrombotic agents | 4 (50.0) |

Unless otherwise indicated, data are mean ± SD values or numbers (%). NIHSS: National Institutes of Health Stroke Scale, BMI: Body Mass Index.
